# Protein folding stress transcriptionally reprograms muscle metabolism

**DOI:** 10.1101/2025.06.20.660805

**Authors:** Jacob Paiano, Mujeeb Qadiri, Suhasini Sharma, Yanhui Hu, Norbert Perrimon

## Abstract

Cellular stress responses crosstalk with many physiological and metabolic pathways. Muscle cells constantly respond to various endogenous stressors while actively maintaining critical metabolic functions for the tissue and whole animal. The molecular mechanisms of how muscle stress responses transcriptionally reprogram metabolic networks are complex and inadequately understood. Using a multi-omics approach of metabolomics, lipidomics, and single-nuclei RNA-sequencing in *Drosophila*, we reconstructed the physiological landscape of muscle during chronic activation of endoplasmic reticulum unfolded protein response (UPR), a stress response that ensures the secretion of vital proteins from muscle, known as myokines. By ectopically expressing a constitutively active form of X-box binding protein 1 (Xbp1), a highly conserved transcription factor (TF) and UPR effector, we found that UPR downregulates key metabolic pathways in muscle, including carbohydrate and purine metabolism, while upregulating a robust lipogenic program enriched for phospholipids and several antioxidant metabolic pathways. Using gene regulatory network (GRN) analysis, we linked these metabolic changes to distinct TF regulon activities. The activation of a single TF, Xbp1, increased the activity of other stress response TFs in muscle, including cap-n-collar (cnc/Nrf2), cryptocephal (crc/Atf4), and sterol regulatory element binding protein (SREBP). Simultaneously, we observed decreased activity of TFs, namely Forkhead box O (FoxO), that resulted in downregulated metabolic pathways critical to muscle function, including oxidative phosphorylation and glycolysis. We propose that these GRNs antagonize each other downstream of UPR to reprogram muscle metabolism away from carbohydrates and towards lipogenesis, offering novel insight into how metabolic rewiring can be transcriptionally controlled in response to chronic tissue damage, even to the detriment of organ function.

## Introduction

Skeletal muscle plays a pivotal role for healthy aging by regulating whole-body metabolism and maintaining mobility (Thyfault et al. 2020; McLeod et al. 2016). Muscle mass declines with age (sarcopenia), as do its stem cells and repair capacity, increasing our susceptibility to chronic diseases (Evans et al. 2010; Kedlian et al. 2024). Elucidating molecular mechanisms of factors that impact muscle growth and health can lead to new therapeutic interventions that promote metabolic healthspan–the number of years living free of disease or disability (Stump et al. 2006; Nishikawa et al. 2021). Chronic metabolic diseases, such as insulin resistance, type 2 diabetes, and obesity, are largely driven by pathological maladaptation of major metabolic organs, such as muscle, fat, and liver (Giangregorio et al. 2024; Barzilai et al. 2012; Lee et al. 2021). Maladaptation to lifestyle (e.g., high calorie diet, sedentation) or genetic factors can predispose muscle to developing metabolic diseases through changing fuel sources, mitochondrial dysfunction, and cellular stress (Amorim et al. 2022; Zhang et al. 2023). Mechanisms of muscle-specific metabolic reprogramming that contribute to metabolic diseases are less understood (Richter-Stretton et al. 2020).

Cellular responses to stress, such as starvation, oxidative radicals, and misfolded proteins, can elicit robust transcriptional programs that alter cell physiology and metabolism until stress is cleared. If stress or damage persists, the chronic activation of stress response programs can drive pathology or aging (Haigis et al. 2010; Kumari 2025; Kourtis et al. 2011). In muscle, a tissue that naturally experiences a great deal of mechanical, nutrient, and oxidative stress, stress responses are deeply interconnected to its physiology and function, with various outcomes for the overall health of the tissue (Deldicque et al. 2012; Tarricone et al. 2006; Lin, et al. 2024; Mensch et al. 2020). In *Drosophila*, we showed that mild mitochondrial dysfunction elicits a mitochondria-specific unfolded protein response that improved muscle health and fly longevity (Owusu-Ansah et al. 2013). We separately showed that increased activity of Forkhead box O (FoxO), a highly conserved transcription factor (TF) activated under conditions of reduced insulin signaling, in muscle limited age-associated protein aggregates, systemically restored proteostasis, and improved fly healthspan and lifespan (Demontis et al. 2010). Our studies, along with many others in *Drosophila* muscle, demonstrate that crosstalk between stress responses and metabolism can significantly affect the health of muscle and the animal (Carney et al. 2023; Zhao et al. 2020; Rai et al. 2021; Green et al. 2018).

Using optogenetics to induce muscle contraction in larvae, mimicking an acute exercise phenotype, we also reported a robust heat shock and protein folding transcriptional response in muscle, suggesting that endoplasmic reticulum (ER) and proteostatic stress are physiological responses to exercise (Ghosh et al. 2022). Exercise has been well documented to improve muscle metabolism and aging (Thyfault et al. 2020; Distefano et al. 2018; Guan et al. 2022). We therefore hypothesized that ER stress responses in muscle may crosstalk with metabolic networks, as others have proposed for ER stress and metabolism in mammalian skeletal muscle (Deldicque et al. 2012). Accumulation of misfolded proteins or increased translational burden in the ER will activate the unfolded protein response (UPR), a highly conserved stress response with three effector branches that transcriptionally upregulate genes to improve protein folding and ER function, such as ER chaperones (Ryoo et al. 2015; Read et al. 2021). One major UPR pathway is the Inositol-requiring enzyme 1 (Ire1) splicing of X box binding protein-1 (Xbp1) mRNA. Unspliced Xbp1 (Xbp1u) does not encode an active TF. Ire1-mediated spliced Xbp1 (Xbp1s) encodes an active TF that upregulates many UPR target genes (Park et al. 2012). Other UPR branches include Pancreatic eIF-2α kinase (PEK) activation of Activating transcription factor 4 (Atf4) and Golgi-mediated cleavage and activation of Activating transcription factor 6 (Atf6). These pathways are largely conserved from flies to humans (Ryoo et al. 2015).

Cellular metabolism is foremost regulated by allosteric interactions between metabolites and enzymes with additional transcriptional regulation by stress and nutrient responsive TFs, such as FoxO (Desvergne et al. 2006, Scholtes et al. 2022). Gene regulatory control of metabolic rewiring in response to homeostatic changes is highly complex and not fully understood at the genome-wide scale (Li et al. 2025). In this study, we performed a multi-omics and systems biology approach to uncover how protein folding stress in muscle can reprogram metabolic networks and impact muscle health. Using *Drosophila* genetic tools to express a constitutively active, spliced Xbp1 protein in adult muscle, we combined single-nuclei RNA-sequencing (snRNA-seq) with mass spectrometry metabolomics and lipidomics to identify all metabolic and gene network changes in muscle downstream of sustained UPR activation. Most strikingly, we observed a robust rewiring away from carbohydrate metabolism and toward lipogenesis, particularly glycerophospholipids. We further found that gene regulatory networks (GRNs) under distinct TFs likely regulate this metabolic reprogramming, demonstrating that chronic stress responses can dramatically alter the physiological landscape of muscle tissue through transcriptional regulation. These metabolic shifts likely represent the increased physiological demand to maintain and expand ER membrane lipids during UPR and, importantly, demonstrate the reallocation of energy resources away from normal physiology and towards organelle repair, even to the potential detriment of the tissue.

## Results

### Chronic muscle unfolded protein response disrupts organismal healthspan and energy homeostasis

To test the effects of sustained UPR activation in muscle, we ectopically expressed in adult muscle using Mhc-GAL4 (myosin heavy chain) a constitutively active, mRNA-spliced form of Xbp1 (i.e., Xbp1s) downstream of the UAS promoter that bypasses endogenous Ire1 regulation (Ryoo et al. 2007). Expression was restricted to the adult stage using temperature-sensitive tubulin-GAL80 and switching newly eclosed flies from 18°C during development to 29°C to permit muscle-GAL4 activity. We chose this strategy of constitutive Xbp1 expression to determine the direct transcriptional consequences of this major arm of the UPR specifically in muscle, instead of relying on damage-induced ER stress, such as tunicamycin feeding, that would lack tissue specificity and activate all UPR branches–and likely other confounding stress responses. We hypothesized that an artificial system of strong Xbp1 activation would permit systems biology analyses that are more robust and easier to interpret than milder, simultaneous activation of several ER and cellular stress networks. To validate that our major findings are applicable to more physiological UPR activation throughout our study, we utilized a mutant form of rhodopsin prone to misfolding in the ER (Rh1^G69D^) under UAS control in adult muscle (Ryoo et al. 2007; Ryoo et al. 2013).

We confirmed the expression and expected TF activity of UAS-Xbp1s using reverse transcription-quantitative polymerase chain reaction (RT-qPCR) on dissected thoraces. We found substantial, and continuous, upregulation of Xbp1s and its major downstream target Hsc70-3/Binding immunoglobulin protein (BiP) at 3d and 7d after switching to 29°C (**Supplemental Figure 1A**). While tracking sustained Xbp1s expression, we found that male and female flies died significantly sooner than controls, on average living only 14 days compared to typical 30+ days (**Figure 1A; Supplemental Figure 1B**). We also recorded climbing ability using negative geotaxis assays during the first ten days of muscle Xbp1s expression and found that climbing scores dramatically decreased over time compared to control flies for both males and females (**Figure 1B; Supplemental Figure 1C**). This suggested that muscle function and/or programmed behavior (i.e., escape response) rapidly declines before fly death during persistent muscle Xbp1 activation.

**Figure 1.**
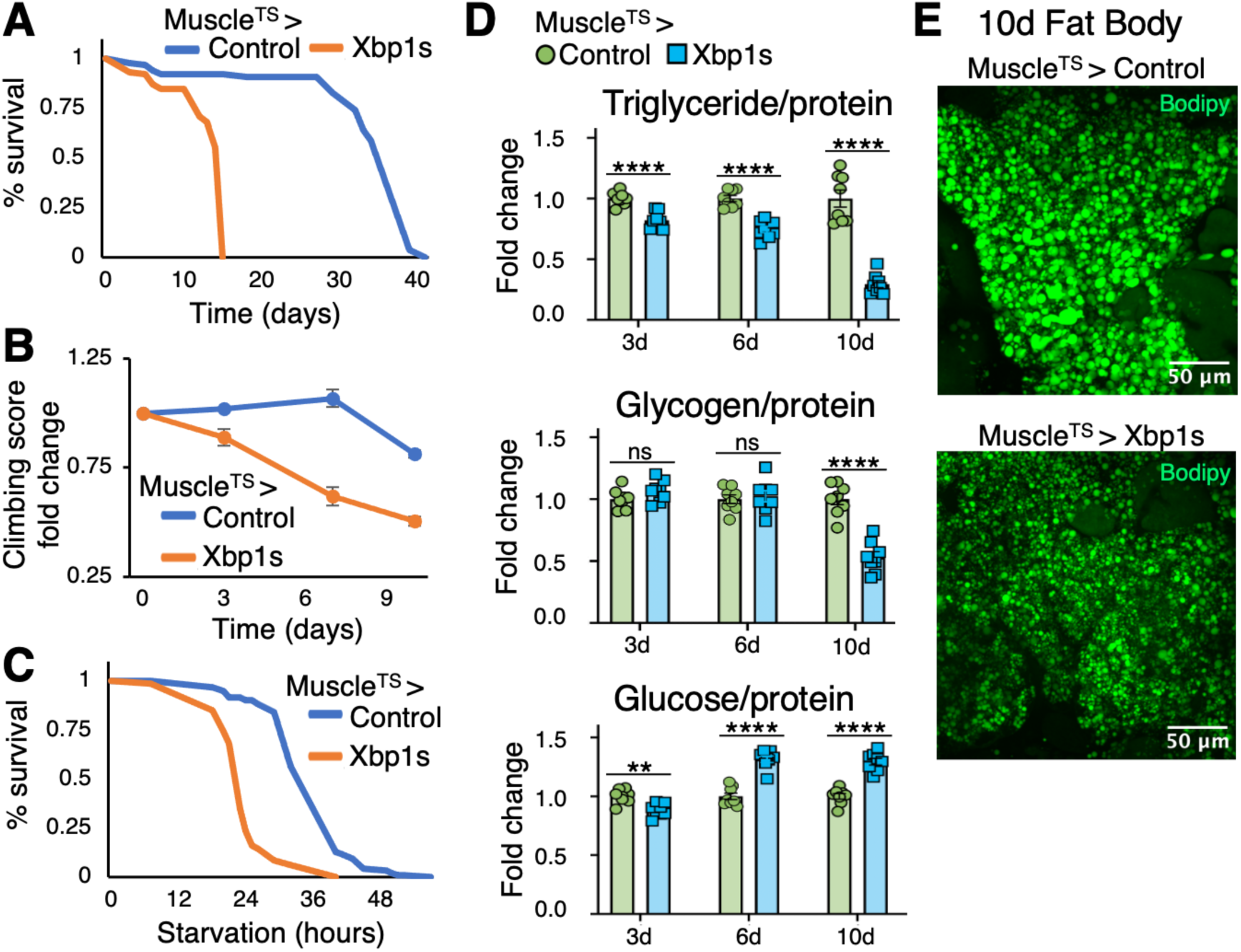
Muscle-specific expression of constitutively active Xbp1 (Xbp1s) decreases fly health/lifespan and disrupts energy homeostasis. **A)** Lifespan of adult male flies placed at 29°C to induce UAS-Xbp1s or UAS-empty vector (control) under temperature-sensitive muscle-GAL4 (Muscle^TS^). **B)** Negative geotaxis climbing score (ranging 1-4) fold change (over 0d time point) in adult males over time. **C)** Wet starvation (1% agar) survival of adult males after 6d Xbp1s muscle expression. **D)** Whole-body detection of triglyceride, glycogen, and glucose, normalized to protein levels, over time in adult males; 1 rep = 8 flies. **E)** Immunohistochemical lipid droplet staining (Bodipy) of abdomen fat body after 10d Xbp1s muscle expression. Significance determined using unpaired T-test.

Because Xbp1s induces ER chaperones, we tested if flies were better able to handle heat shock stress. We expressed Xbp1s in muscle for 6 days at 29°C then heat-shocked flies at 39°C while frequently recording fly death counts. While control flies remained active for 30-40 minutes, Xbp1s-expressing male and female flies died significantly sooner, especially females, indicating that prolonged muscle-specific Xbp1 activation increases organismal sensitivity to heat shock despite upregulating protective proteostasis genes (**Supplemental Figure 1D**). Climbing defects and heat shock sensitivity suggested a general decline in fly healthspan.

We next tested if flies were similarly sensitive to starvation by transferring flies from standard food to 1% agar after 10 days of Xbp1s expression in muscle. As with heat shock, we found Xbp1 activation in muscle significantly decreases survival of both male and female flies during wet starvation (**Figure 1C; Supplemental Figure 2A**). Increased starvation sensitivity prompted us to measure macronutrient storage. We found that whole-body triglyceride (TG) and glycogen storage substantially decrease over time, alongside a modest, yet consistent, rise in glucose (**Figure 1D**). To validate that these whole-animal phenotypes were not due to GAL4 expression in non-muscle tissues, we confirmed TG depletion using two separate muscle-specific GAL4 drivers, dMef2 (pan-muscle) and Act88F (indirect flight muscle), and found reductions in whole-fly TG in both males and females (**Supplemental Figure 2B**).

We then asked whether this result was specific to Xbp1 TF activity or more generalizable to physiological muscle UPR induction. To do so, we expressed Rh1^G69D^ (a misfolding-prone mutant of rhodopsin) in adult muscle. We first confirmed that Rh1^G69D^ expression induced endogenous Ire1-mediated Xbp1 splicing and Hsc70-3/BiP upregulation in thoraces by RT-qPCR, demonstrating its capacity to sustain UPR in muscle via continuous protein misfolding, although not as strongly as UAS-Xbp1s (**Supplemental Figure 2C**). We next found that, like Xbp1s, whole-fly TG was reduced >50% while glucose levels rose slightly after chronic Rh1^G69D^ expression (**Supplemental Figure 2D**). The substantial decrease in TG suggests communication between muscle and fat body, the major tissue of TG storage, during UPR activation. Consistent with this, we observed decreased lipid droplet size in abdomen fat body after 10 days of Xbp1 activation in muscle (**Figure 1E**). Altogether, these results indicate that prolonged, muscle-specific protein folding stress diminishes healthspan and lifespan and disrupts muscle and organismal metabolism through unknown mechanisms.

### Characterizing the transcriptome of Xbp1-activated muscle using single-nuclei RNA-sequencing

To uncover the systemic pathways underlying health and metabolic decline during muscle UPR activation, we performed whole-body (heads removed) snRNA-seq on male flies after 3 days of Xbp1s expression in muscle. We chose this early time point to capture how Xbp1 initially reprograms the muscle transcriptome before healthspan and metabolism severely disrupted, avoiding overly complex phenotypes (**Figure 1**). As expected with our whole-body snRNA-seq protocol (Liu et al. 2025), we found significant representation of all major body tissues in the snRNA-seq, with clusters representing indirect flight muscle (hereby referred to as “flight muscle”; analogous to mitochondria-rich, slow twitch (type I-like) muscle fibers (Bryantsev et al 2019)), “other muscle” (likely heterogeneous population including jump muscle, analogous to fast twitch (type II-like) muscle fibers (Bryantsev et al 2019)), fat body (adipose and liver-like functions), oenocytes (liver-like functions), hemocytes (phagocytic innate immune cells), epithelium, peripheral neurons, and ventral neurons (**Figure 2A; Supplemental Figure 3A**). The pan-muscle Mhc-Gal4 driver should strongly upregulate Xbp1/Xbp1s in both flight muscle and other muscle, which we observed (**Figure 2B**). Alongside Xbp1 upregulation, we found increased expression of other major UPR effectors, such as Atf6 and cryptocephal (crc)/Atf4 (Ryoo 2015), and known UPR target chaperones and protein folding genes (**Figure 2C**). We also saw upregulation of various genes involved in protein targeting and ER export, suggesting that Xbp1s activation does not downregulate ER trafficking and protein secretion (**Figure 2C**).

**Figure 2.**
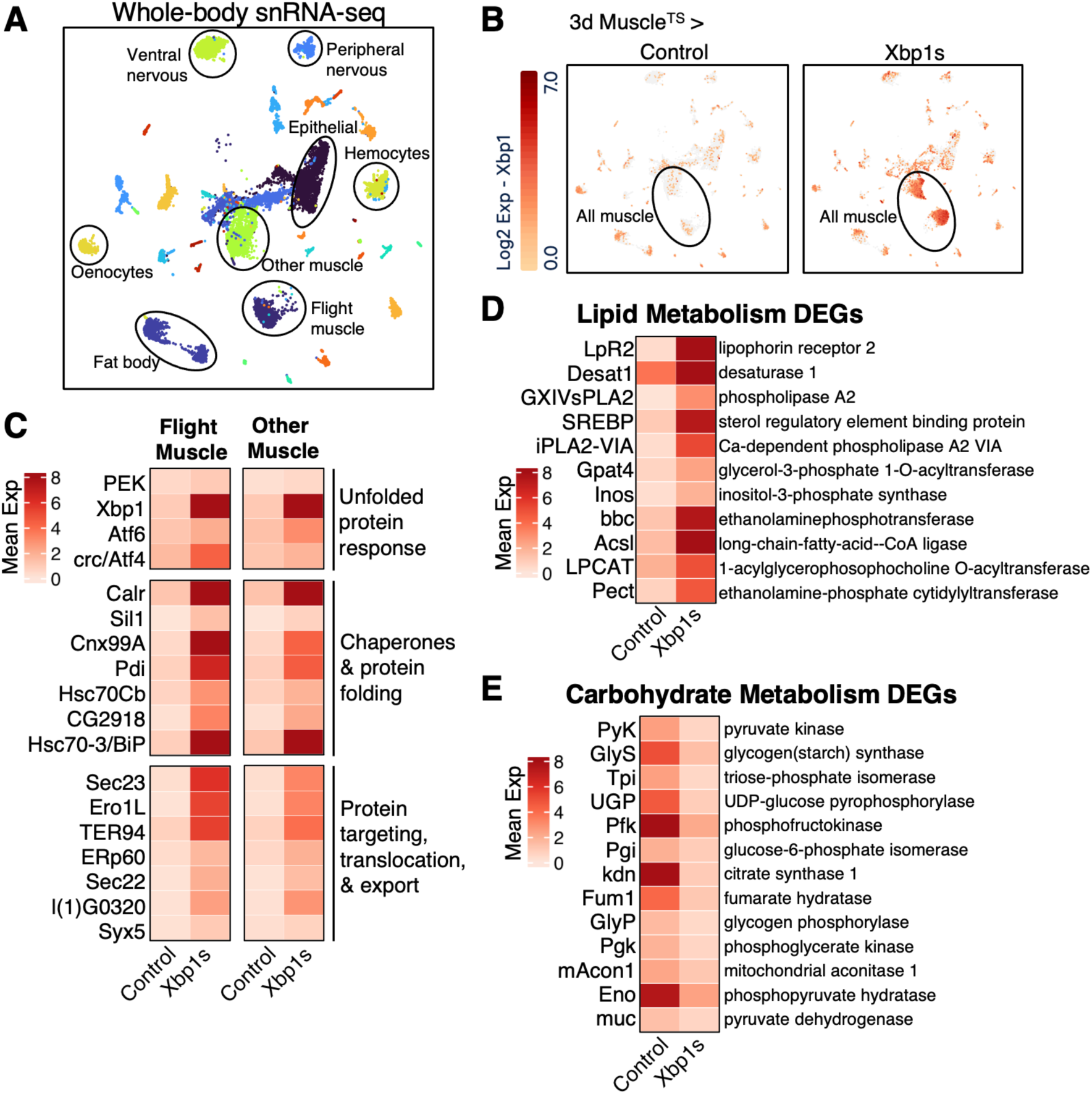
Single-nuclei RNA-sequencing (snRNA-seq) confirmed muscle-specific upregulation of Xbp1 and UPR, while revealing increased lipid and decreased carbohydrate metabolic genes. **A)** UMAP of whole-body (heads removed) snRNA-seq clusters representing major tissues, including two muscle clusters annotated as “flight muscle” and “other muscle”. **B)** UMAP confirming muscle-specific upregulation of Xbp1 mRNA after 3d Xbp1s expression. **C)** Heatmap of flight muscle and other muscle cluster mean expression for various UPR effectors, chaperones, and protein export genes, confirming the expected transcription factor activity of Xbp1s. **D, E)** Differentially expressed gene (DEG) analysis of flight muscle identified many significantly upregulated lipid metabolism genes **(D)** and downregulated carbohydrate metabolism genes **(E)**, suggesting a transcriptionally controlled metabolic reprogramming away from glycogen/glucose and towards fatty acid metabolism.

As Xbp1s is a constitutively active TF, we expected many changes in transcriptional programs after 3-day expression in muscle. Indeed, we found substantial shifts in UMAP cluster projections for both flight muscle and other muscle between control and Xbp1s flies, indicating reprogramming of many genes, likely beyond UPR, in muscle transcriptomes (**Supplemental Figure 3B**). Importantly, we did not see UMAP cluster shifts for other tissues, confirming the restriction of Xbp1s reprogramming to muscle (**Supplemental Figure 3B**). To interrogate these transcriptomic changes, we determined all differentially expressed genes (DEGs) for Xbp1s versus control. Filtering by log2FoldChange > 1.0 and percent expression >10% of Xbp1s cells, we identified 393 significantly upregulated DEGs in Xbp1s flight muscle. Using the PAthway, Network, and Gene-set Enrichment Analysis (PANGEA) data analysis tool (Hu et al 2023), we performed overrepresentation analysis on these DEGs for enriched Gene Ontology (GO) slim subset (SLIM2) biological process (BP) *Drosophila* terms. Top terms included expected ER processes, such as protein secretion, protein folding, and endomembrane system organization (**Supplemental Figure 4A**). We additionally found response to stress, autophagy, cell redox homeostasis, lipid transport, and various muscle-related enriched terms. We next created gene set node graphs to visualize shared genes between UPR and response to stress terms and identified many stress response genes outside of ER function that are upregulated, including several critical metabolic regulators such as sterol regulatory element binding protein (SREBP), Thor/4E-BP, Autophagy related-1 (Atg1), and Imaginal morphogenesis protein-Late 2 (Impl2) (**Supplemental Figure 4B**).

To explore these metabolic changes further, we identified upregulated Kyoto Encyclopedia of Genes and Genomes (KEGG) *Drosophila* terms in flight muscle. In addition to protein export and protein processing in ER, several lipid metabolic processes were most enriched, including phosphatidylethanolamine biosynthesis, glycerophospholipid, inositol phosphate, ether lipid, and arachidonic acid metabolism (**Supplemental Figure 4C**). This was consistent with literature describing the close interplay of ER stress and phospholipid synthesis to expand ER membrane (Fagone et al 2009, Sriburi et al 2004, Sriburi et al 2007). Non-lipid metabolic terms, including protein catabolism, glutathione, cofactors, and various amino acid terms were also enriched. To determine if these metabolic shifts were part of Xbp1s-induced stress response programs, we generated gene set node graphs to visualize the interconnectedness of enriched stress and metabolic terms. Indeed, we found many shared gene nodes between metabolic pathways, “response to stress”, and “protein processing in ER” networks, suggesting these metabolic changes are specific to the programmed stress-induced needs of the tissue (**Supplemental Figure 4D**). Importantly, many of the gene nodes represent fly orthologues of rate-limiting metabolic enzymes (Zhao et al. 2009), such desaturase 1 (Desat1), acyl-CoA synthetase long-chain (Acsl), lysophosphatidylcholine acyltransferase (LPCAT), glycerol-3-phosphate acyltransferase 4 (Gpat4), glutamine synthetase 2 (Gs2), and nicotinamide mononucleotide adenylyltransferase (Nmnat) (**Supplemental Figure 4D**).

We next interrogated downregulated transcriptional programs between control and Xbp1s flight muscle. We filtered DEGs for downregulated genes with log2FoldChange < -1.0 and percent expression >10% in control cells and performed the same GO and KEGG term overrepresentation analyses on these 278 downregulated DEGs. Significantly downregulated GO terms included actin cytoskeleton, muscle structure development, locomotory behavior, and circadian rhythm (**Supplemental Figure 5A**). These pathways shared many gene nodes (**Supplemental Figure 5B)**, suggesting a programmed coordination in decreased expression of these networks that may contribute to our observed climbing defects and declined muscle function (**Figure 1B**, **Supplemental Figure 1C**).

We also found strong enrichment of the GO terms, “generation of precursor metabolites and energy” and “carbohydrate metabolic process”. KEGG enrichment further identified tens of downregulated pathways and reactions related to energy and carbohydrate metabolism, including glycogen metabolism, citrate/TCA cycle, pentose phosphate pathway, oxidative phosphorylation, and mitochondrial complexes (**Supplemental Figure 5C**). Individual pathways were highly interconnected and comprehensive, indicating a global decrease in the expression of genes related to carbohydrate processes, mitochondrial respiration, and purine metabolism (**Supplemental Figure 5D**). Critically, downregulated metabolic nodes included many rate-limiting enzymes, such as glycogen synthase (GlyS), glycogen phosphorylase (GlyP), phosphofructokinase (Pfk), pyruvate kinase (PyK), and pyruvate dehydrogenases (Pdha, muc) (**Supplementary Figure 5D**). Downregulated oxidative phosphorylation nodes included F-type ATPases, cytochrome bc1 complex, and NADH dehydrogenase beta and alpha subcomplexes (**Supplemental Figure 5D**).

Altogether, our DEG pathway analyses strongly suggested that 3 days of Xbp1 activation in flight muscle substantially rewired metabolism through transcriptional control of many lipid and carbohydrate processes; generally shifting cells away from carbohydrate consumption and oxidative phosphorylation and towards lipogenesis via upregulation of key lipid metabolic genes, including fatty acid uptake lipophorin receptor 2 (LpR2), and downregulation of key carbohydrate metabolic genes (**Figure 2D, 2E**). We also performed overrepresentation analyses on the 226 upregulated DEGs and 154 downregulated DEGs in Xbp1s-expressing “other muscle” cells. We identified similarly upregulated lipogenesis KEGG terms and rate-limiting enzymes, including fatty acid biosynthesis via fatty acid synthase 1 (FASN1) and acetyl-CoA carboxylase (ACC) (**Supplemental Figure 6A, 6B**). Consistent with flight muscle, we found tens of downregulated KEGG terms related to carbohydrate metabolism and oxidative phosphorylation, with many of the same downregulated rate-limiting enzymes (**Supplemental Figure 6C, 6D**). This suggested that Xbp1 has the capacity to rewire energy programs similarly across muscle tissues, regardless of type I-like or type II-like muscle fibers. These analyses also suggest that robust activation of stress response TF effectors, like Xbp1, can transcriptionally reprogram metabolic tissues to rewire specific energy demands.

### Characterizing the metabolome and lipidome of Xbp1-activated muscle

Our snRNA-seq analysis of upregulated and downregulated DEGs identified many altered transcriptional programs, including both expected UPR and stress signatures and unexpected metabolic network rewiring. Regulation of metabolic pathways is complex, and changes in gene expression are only predictive of potential metabolic outcomes, as allosteric regulation of enzymes is most utilized by cells. To investigate our findings that lipogenic programs increased and carbohydrate programs decreased, we performed liquid chromatography-mass spectrometry (LC-MS) polar metabolomics and non-polar lipidomics on dissected adult, male thoraces with or without 6 days of Xbp1s muscle expression. While our snRNA-seq was collected at 3 days, we chose to profile metabolomic and lipidomic changes at 6 days, a slightly later time point in which flies begin to climb poorly (**Figure 1B**), are slightly hyperglycemic, but just before dramatic disruptions in whole-body energy homeostasis (i.e., decreased TG and glycogen storage) are observed (**Figure 1D**). We hypothesized that this time point would reveal metabolomic and lipidomic changes that may contribute to muscle dysfunction, eventual organismal metabolic dysfunction, and are direct consequences of early transcriptional reprogramming by Xbp1s in muscle.

LC-MS polar metabolomics showed that many thorax metabolites increased or decreased after 6 days of Xbp1s expression (**Figure 3A, Supplemental Figure 7A**). The most apparent metabolite increases were associated with protective oxidative stress responses, including citrulline and ornithine, products of arginine, proline, and glutamate metabolism (**Figure 3A, Supplemental Figure 7A**). Arginine, citrulline, and ornithine have been extensively studied in the context of muscle physiology, exercise, and aging. Studies have suggested they promote human growth hormone-mediated muscle building and increased exercise capacity via nitric oxide synthesis (McConnell 2007; Viribay et al 2020; Gough et al 2021; Demura et al 2010). In fact, all three are commercially available as sports supplements. Additionally, spermidine, a polyamine generated from ornithine, significantly increased (**Figure 3A, Supplemental Figure 7A**). Spermidine, along with other polyamines, has been suggested to promote autophagy and mitophagy in skeletal muscle, combat aging, and promote muscle growth (Galasso et al 2023; Lee et al 2011). There was also a significant increase in uric acid and allantoin after Xbp1 activation, metabolic signatures of scavenging oxidative species in skeletal muscle post-exercise (**Figure 3A, Supplemental Figure 7A**) (Hellsten et al 2001; Kand’ar et al 2008). Together, these increased metabolites suggested a protective metabolic response to oxidative stress that may be beneficial for muscle repair after damage and proteostatic stress.

**Figure 3.**
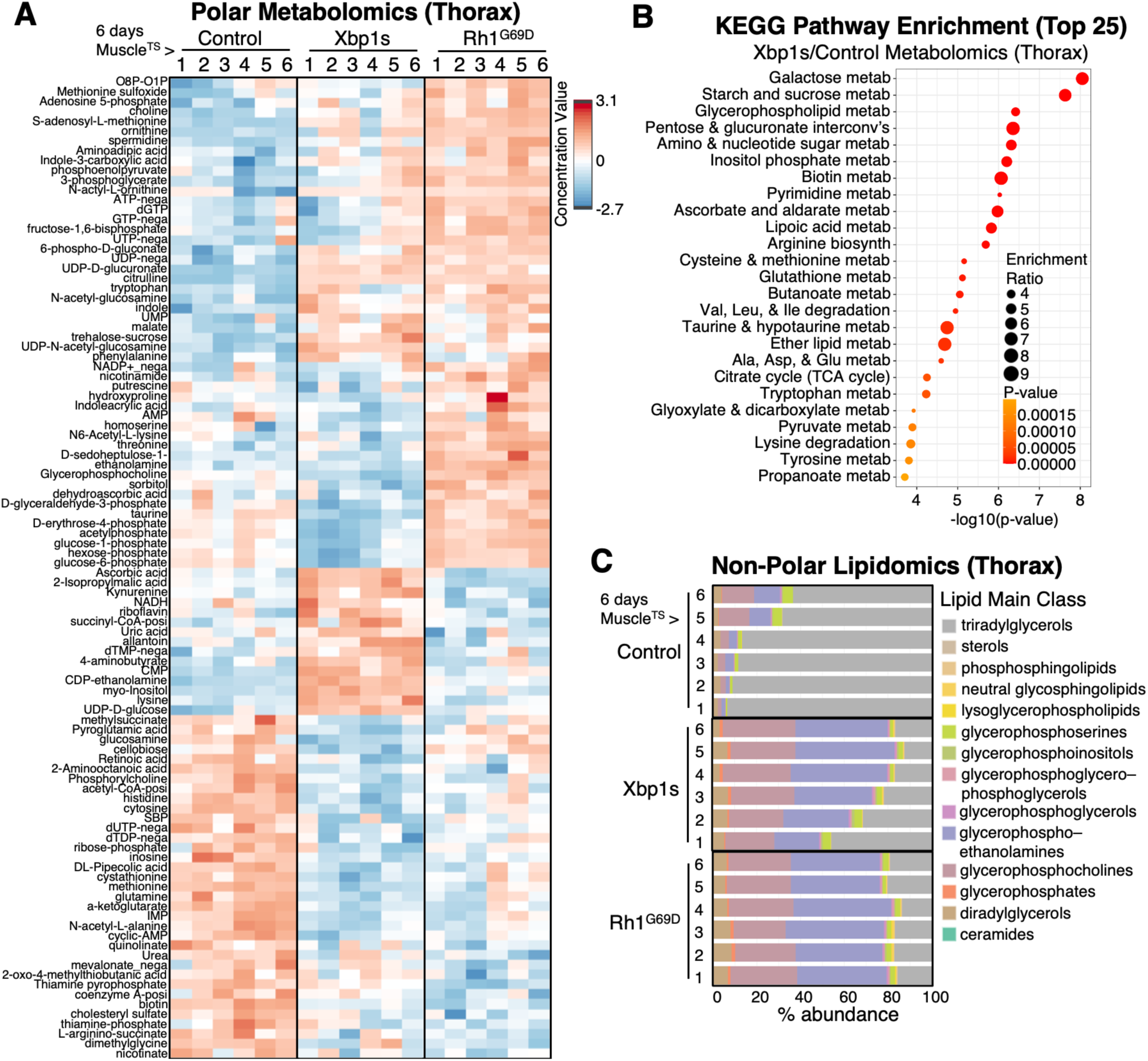
Mass spectrometry metabolomics and lipidomics of UPR active thoraces showed altered carbohydrate, phospholipid, and antioxidant pathways. **A)** Polar metabolomics of dissected thoraces after 6d Xbp1s or misfolded rhodopsin (Rh1^G69D^) muscle expression; 1 rep = 25 thoraces. Heatmap of top 100 most significantly changed metabolite abundances. **B)** Most enriched annotated KEGG terms in list of changed polar metabolites from Xbp1s versus control thoraces. Enrichment indicates pathway disruption compared to control, not necessarily increased or decreased abundance. **C)** Nonpolar lipidomics from the sample thorax samples as **(A)**. Xbp1s and Rh1^G69D^ expression greatly increased the relative abundance of several main classes of phospholipid related species compared to controls.

To understand if metabolic changes were specific to Xbp1s expression or more generalizable to physiological ER stress, we also performed metabolomics on muscle-expressing Rh1^G69D^ thoraces. Principle component analysis (PCA) of six biological replicates (25 thoraces per replicate) of control, Xbp1s, and Rh1^G69D^ showed that the largest variance (21.9%) in the data was between control and Xbp1s/Rh1^G69D^, while the second largest (16.9%) was between Xbp1s and Rh1^G69D^, indicating that Xbp1s and Rh1^G69D^ alters muscle metabolome largely similarly but with some significant differences (**Supplemental Figure 7B**). Of the top 100 most altered metabolites, virtually all decreased metabolites were shared between Xbp1s and Rh1^G69D^ (**Figure 3A, Supplemental Figure 7A, 7C**). Many increased metabolites were also shared, although, several were unique to either Xbp1s or Rh1^G69D^ (**Figure 3A**). To further interrogate these differences, we used MetaboAnalyst 6.0 quantitative enrichment analysis (Pang et al. 2024) to determine significantly altered KEGG metabolite pathways. Despite some differences in increased metabolite abundances, most dysregulated pathways were shared between Xbp1s and Rh1^G69D^ (**Figure 3B, Supplemental Figure 7D**). This may have reflected metabolite differences within specific reactions of large KEGG terms, the additional activation of non-Ire1/Xbp1 UPR branches downstream of Rh1^G69D^, or that Rh1^G69D^ induces endogenous Xbp1 less strongly than UAS-Xbp1s (**Supplemental Figure 2C**). Overall, 6 days of Xbp1/UPR activation in muscle significantly alters muscle metabolomic networks.

KEGG set enrichment analysis revealed several disrupted amino acid pathways, such as arginine, which is consistent with increases in citrulline and ornithine production. Cysteine, methionine, and glutathione pathways were also altered (**Figure 3B, Supplemental Figure 7D**). Methionine significantly decreased, while S-adenosylmethionine (SAM) and S-adenosylhomocysteine (SAH) increased, suggesting activation of the methionine cycle and increased glutathione cycle, which is a critical redox pathway to limit oxidative stress (**Figure 3A**) (Diaz-Vivancos et al. 2015). Moreover, methionine itself can directly scavenge free radicals to form methionine sulfoxide, of which we also observed increased abundance (**Figure 3A**) (Luo et al. 2009; Pamplona et al. 2006). These findings further support a change in muscle metabolome to induce an antioxidant program downstream of protein folding stress.

Several carbohydrate metabolic pathways were also disrupted, such as citrate/TCA cycle, glycolysis and gluconeogenesis, pentose phosphate pathway, and pyruvate metabolism (**Figure 3B, Supplemental Figure 7D**), consistent with our snRNA-seq analysis that showed decreased expression of genes and enzymes related to these pathways (**Figure 2E, Supplemental Figure 5C, 5D**). After Xbp1 activation, we observed decreased glucose-1-phosphate and glucose-6-phosphate, byproducts of glycogen breakdown (**Figure 3A, Supplemental Figure 7A**). We also found increased UDP-glucose (converted from glucose-1-phosphate), a building block for the synthesis of sucrose, polysaccharides, and other metabolites, such as UDP-glucuronate, which we also found to be increased. Altogether, these findings suggest a significant dysregulation in glycogen and glucose metabolism and support our transcriptomic findings that carbohydrate metabolism is substantially disrupted in muscle downstream of UPR activation.

In addition to carbohydrates, several lipid metabolic pathways were also enriched in both Xbp1s and Rh1^G69D^ thoraces, including glycerophospholipid, ether lipid, and lipoic acid metabolism (**Figure 3B, Supplemental Figure 7D**). Metabolomics captured certain polar compounds, such as enrichment of ethanolamine, CDP-ethanolamine, phosphocholine, and glycerophosphocholine, key metabolites for glycerophospholipid generation (**Figure 3A, Supplemental Figure 7A, 6B**). These metabolites supported our snRNA-seq analysis that showed increased lipogenic transcriptional programs, such as glycerophospholipid and ether lipid metabolism (**Figure 2D, Supplemental Figure 4C, 4D**). To fully characterize changes to muscle lipidome, we performed nonpolar lipidomics on the same thorax samples as polar metabolomics to capture all changes in lipid species and used LipidSig 2.0 for all downstream analysis (Lin et al 2021; Liu et al 2024). PCA revealed significant differences between control and UPR-activated thoraces; however, unlike metabolomics in which some increased polar metabolites differed, Xbp1s- and Rh1^G69D^-expressing thoraces clustered together, indicating that the lipidomic profiles were virtually identical in composition (**Supplemental Figure 8A**). Overall, UPR-activated thoraces exhibited substantially increased normalized lipid counts, averaging 20-to-30-fold increases over control, thus confirming robust lipogenesis (**Supplemental Figure 8B**).

Consistent with metabolomics and transcriptomics, 50-80% of all detected lipids in UPR-activated thoraces were comprised of main lipid classes related to phospholipids, including glycerophosphoserines (PS), glycerophosphoinositols (PI), glycerophosphoglycerols (PG), glycerophosphoethanolamines (PE), glycerophosphocholines (PC), and glycerophosphates (PA) (**Figure 3C**). These increased classes were largely comprised of 1,2-diacyl (∼25%) or 1-alkyl,2-acyl (∼60%) glycerophospholipids, with a small fraction of ceramide phosphocholines/sphingomyelins (∼10%) (**Supplemental Figure 8C**). We next asked which specific lipid reactions were affected that cause these changes in class composition after UPR induction. We first identified all differentially abundant lipid species followed by network analysis using LipidSig. Pathway activity network and lipid reaction network analyses were highly enriched for conversion of diacylglyercol (DG) to various glycerophospholipids, mostly PE, PC, PG, PS, and PI, with further conversion to lysophospholipids, such as LPE, LPS, and LPG (**Supplemental Figure 9A, 9B**). Network analyses also revealed significant conversion of akyl ether linkage DG (DG O-) to ether liked PE (PE O-) and PC (PC O-) and the conversion of ceramide to hexosylceramide, a type of sphingolipid with attached sugar molecule. Altogether, our lipidomics analysis revealed the extent to which ER stress reprograms muscle physiology to increase glycerophospholipid, ether lipid, and sphingolipid metabolism.

### Reconstructing Xbp1-induced gene regulatory networks

snRNA-seq DEG analyses predicted altered metabolic pathways that were highly interconnected with Xbp1s-induced stress responses (**Supplemental Figure 4D, Supplemental Figure 5D**), which we then confirmed with LC-MS metabolomic and lipidomic profiling. The strong predictability of metabolic outcomes from the transcriptomics prompted us to hypothesize that we could reconstruct GRNs downstream of Xbp1 activation that regulate our observed metabolic changes to provide more mechanistic insight into how ER stress transcriptionally regulates muscle metabolism.

To interrogate GRNs in our muscle snRNA-seq clusters, we first utilized Single-Cell rEgulatory Network Inference and Clustering (SCENIC) to characterize all TF networks and their target genes after Xbp1 activation (Aibar et al. 2018). Briefly, SCENIC works by building co-expression modules of TFs and potential co-regulated gene targets. It then refines these modules using TF binding/motif databases to construct higher confidence TF “regulons” (i.e., co-regulated gene modules predicted to be controlled by a single TF). Finally, it uses the tool AUCell to score regulon enrichment in the top percentile of expressed genes per cell. This score can be interpreted as a TF regulon activity score for each cell in the snRNA-seq dataset, which can then be averaged across a cluster to predict TF activity for a given tissue. Because of inherent stochasticity in SCENIC that cause slight variations in regulon target genes each run, we ran SCENIC twenty times on Xbp1s flight muscle cells and filtered for only regulons and target genes present in >80% of runs. We then scored each cell using AUCell for the filtered regulons and target genes to obtain a list of active TFs and their predicted gene regulatory activity scores downstream of Xbp1 activation. We then used AUCell to score these Xbp1 regulons in control flight muscle cells to determine which TFs change their activity between control and Xbp1 activation (**Supplemental Figure 10A**).

We further filtered regulons by AUCell scores greater than 0.1 in either control or Xbp1s cells to focus on the most impactful and active GRNs, as regulons scored <0.1 were generally lowly expressed TFs (not shown). We additionally discarded any regulon not technically annotated as a TF, as many regulons were constructed around non-TF nucleic acid binding proteins, such a chromatin remodelers. Finally, we calculated the regulon activity score fold change for Xbp1s/control and kept only regulons with increased activity fold change >1.2 or decreased fold change <0.8. This produced a list of highest confidence and most impactful TF network changes between control and Xbp1s flight muscle, including 8 increased and 9 decreased regulons (**Supplemental Figure 10B**). As expected, Xbp1 activity itself increased and included 204 predicted target genes that were enriched for KEGG and GO terms similar to flight muscle upregulated DEGs, including protein processing in ER, protein folding and export, glycerophospholipid metabolism, autophagy and mitophagy, and lipid transport (**Supplemental Figure 10C**).

Interestingly, other increased regulons included major stress and metabolic TFs, such as SREBP, crc/ATF4, Cyclic-AMP response element binding protein B (CrebB), Mondo/ChREBP (Carbohydrate Response Element Binding Protein), and the antioxidant regulator cap-n-collar (cnc)/Nuclear factor erythroid 2-related factor 2 (NRF2), suggesting Xbp1 may coordinate with these TFs for ER stress-associated metabolic rewiring (**Supplemental Figure 10C**). There were also several Xbp1s regulons with higher scores in control flight muscle, suggesting a downregulation of their activity, including other important stress and metabolism regulators, such as the mechanistic Target of rapamycin (mTOR) controlled FoxO, Forkhead box K (FoxK), the insect nuclear hormone receptor Ecdysone Receptor (EcR), mirror (mirr)/Iroquois, and the hypoxia responder similar (sima)/hypoxia-inducible factor 1 alpha (HIF1A) (**Supplemental Figure 10C**).

Having identified potential GRNs in flight muscle, we then performed the same SCENIC analysis on “other muscle” cells, first building Xbp1s-associated regulons and AUCell scoring in both control and Xbp1s cells (**Supplemental Figure 10D**). We found only six TFs with predicted increased activity and no TFs with decreased activity in Xbp1-activated other muscle cells (**Supplemental Figure 10E**). Of the increased regulons, Xbp1, SREBP, and cnc were shared. Additional stress regulators such as Cyclic-AMP response element binding protein A (CrebA) and Activating transcription factor 3 (Atf3) were also increased. Overall, both flight muscle and other muscle regulon analyses identified several potential TFs that coordinated increased stress and metabolic programs; however, neither were very successful at robustly identifying TFs with decreased activity that may control downregulated programs.

To address this, we devised another SCENIC strategy to specifically predict differentially active regulons between control and Xbp1s cells. This strategy employed the same SCENIC pipeline described above but with the exception that we included both control and Xbp1s cells together as input for TF-target genes co-expression. We hypothesized that this approach would better assign downregulated genes to TFs with decreased activity, as including control cells would help build regulons that include co-expression of TFs and targets in normal muscle that can be scored as downregulated in Xbp1s cells. Applying the same filtering and fold change cutoffs as described, we indeed identified more regulons with greater increased fold change and lower decreased fold change in activity, giving us higher confidence to predict altered GRNs (**Figure 4A, 4B**). All increased regulons from our initial analysis were present, as were many of the decreased, such as FoxO, FoxK, mirr, and sima, indicating our strategy to include both control and Xbp1s cells only improved our ability to detect differentially active regulons and did not hamper it. We also used this strategy to determine differentially active regulons in other muscle cells and obtained similar results with more regulons having increased or decreased fold changes and identified several predicted downregulated TFs (**Supplemental Figure 11A, 11B**).

**Figure 4.**
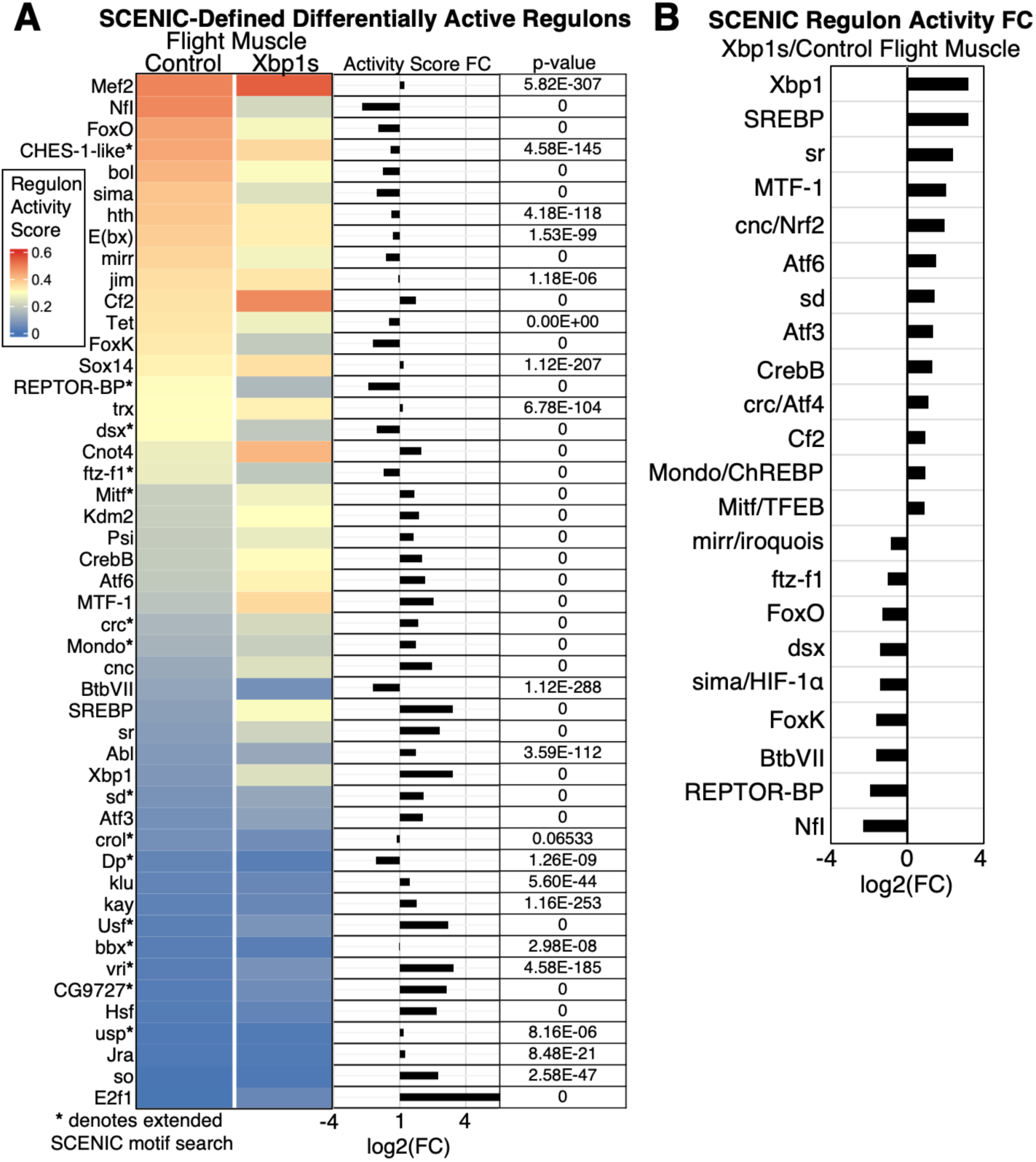
Single-Cell rEgulatory Network Inference and Clustering (SCENIC) identification of differentially active transcription factor regulons between Xbp1s and control snRNA-seq. **A)** Heatmap of mean regulon activity scores between control and Xbp1s flight muscle cells with fold change and significance. For some regulons, SCENIC required an extended database motif search to finish construction (denoted with *). **B)** Finalized list of regulons and fold change from **(A)** after removing non-transcription factor regulons and filtering for >0.1 activity score and either >1.2 or <0.8 fold change for Xbp1s/control. Xbp1 itself was predicted to be the most increased regulon in flight muscle, confirming the accuracy of our SCENIC pipeline.

### Assigning metabolomic and lipidomic changes to Xbp1-induced GRNs

We next sought to mechanistically link altered GRNs to metabolomic and lipidomic rewiring and devised a strategy to assign changes in metabolites first to DEGs, then to differentially active TF regulons (**Figure 5A**). To identify relevant DEGs, we relied on metabolic databases with annotated *Drosophila* genes known for each pathway, such as the enriched KEGG fly terms we identified in metabolomics (**Figure 3B**). We began with all KEGG terms (43 total) from MetaboAnalyst that were significantly enriched in altered metabolite abundances between control and Xbp1s-expressing thoraces. To expand our terms and gene annotations, we next manually selected similar pathways (97 total) from the BioCyc Genome Database Collection (Karp et al 2019). For lipidomics, we selected KEGG and BioCyc terms (13 total) that included any significantly changed lipid class, such as glycerophospholipids and sphingolipids (**Figure 3C, Supplemental Figure 8C**). We additionally included results from LipidSig pathway activity network and lipid reaction network analyses (**Supplemental Figure 9A, 9B**), as LipidSig annotates each lipid reaction with associated human genes. We consolidated individual reactions to a final list of terms (156 reactions total) and mapped all high-confidence *Drosophila* orthologues from annotated human genes to create a fly-specific term for each lipid reaction.

**Figure 5.**
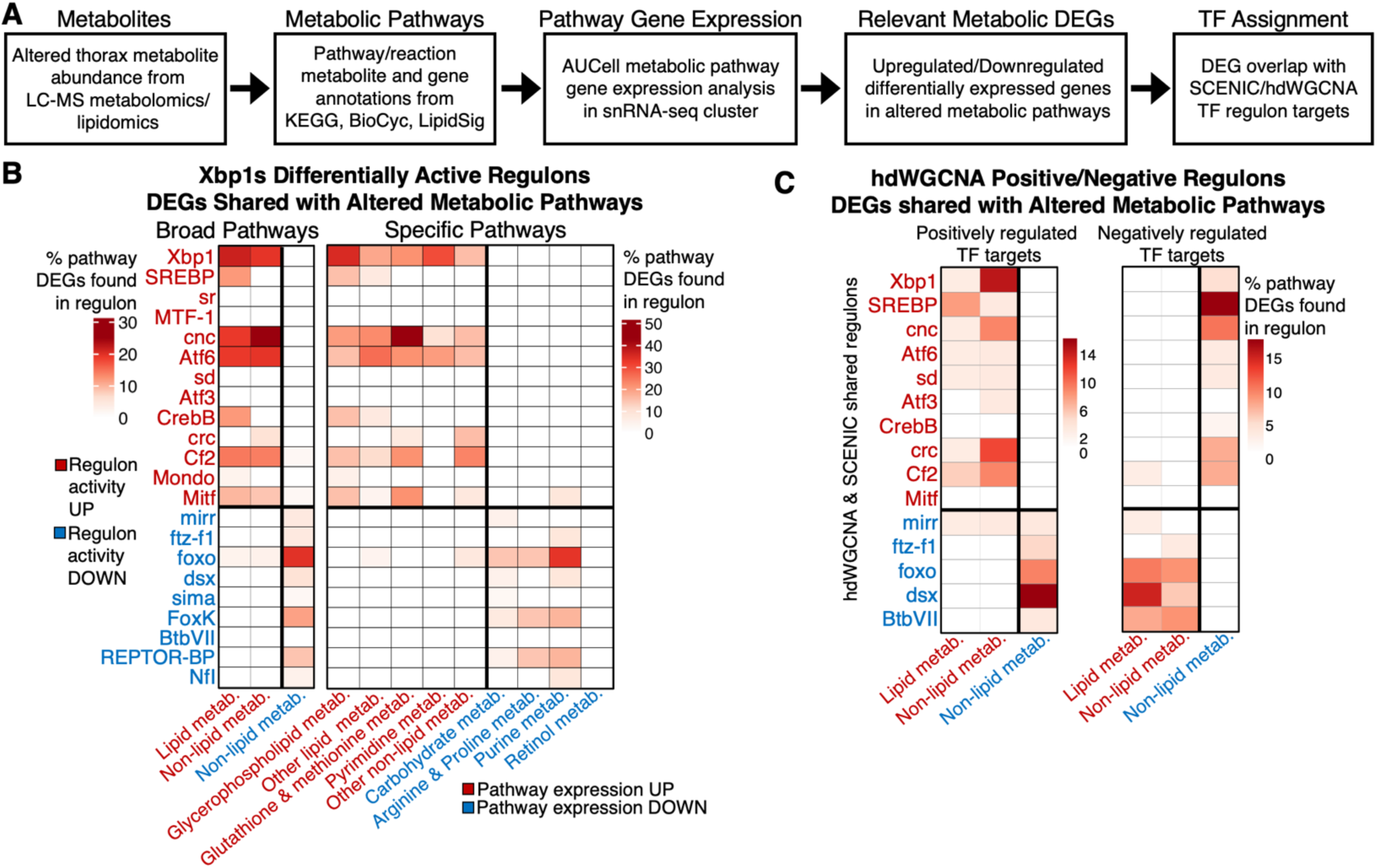
Integration of metabolomics and lipidomics with snRNA-seq to predict gene regulatory networks driving metabolic network changes in Xbp1 activated flight muscle. **A)** Computational workflow starting with empirical changes in polar and nonpolar metabolite abundances to identify disrupted metabolic pathways. Pathways were then scored with AUCell to determine changes in gene expression (i.e., potential network-level transcriptional control of pathway). Annotated genes in those pathways that were also significant DEGs between Xbp1s and control flight muscle were then intersected with SCENIC- or high dimensional weighted gene co-expression network analysis (hdWGCNA)-defined regulons to predict TFs that regulate metabolic changes. **B)** Heatmap depicting the percentage of DEGs in a given pathway (categorized into either broad pathways or specific pathways) that were also found in a given SCENIC differentially active regulon. Regulons with increased activity (in red) were enriched for DEGs found in upregulated lipid and non-lipid metabolic pathways (in red). Similarly, regulons with decreased activity (in blue) were enriched for DEGs found in downregulated non-lipid metabolic pathways (in blue). **C)** Same percent DEG analysis as in “**B**” but with hdWGCNA regulons divided into predicted positively regulated or negatively regulated target genes per TF. Only regulons shared between SCENIC and hdWGCNA are shown. Generally, regulons with increased activity were predicted to positively regulate upregulated DEGs and negatively regulate downregulated DEGs, and vice-versa for regulons with decreased activity.

With the goal to identify GRNs that regulate altered metabolic networks, we acknowledged that not all metabolic pathways are likely to be transcriptionally controlled at the network level. To focus only on annotated pathways that represent network-level transcriptional regulation, we next performed AUCell scoring of each pathway (140 metabolomic and 169 lipidomic terms) for snRNA-seq Xbp1s and control flight muscle cells (**Supplemental Figure 12A-C**). We reasoned that increased or decreased AUCell scores would indicate pathways that change gene expression at a network level and may be controlled by specific GRNs. Using the same cutoffs as our SCENIC filtering, we kept only pathways with Xbp1s/control fold change in AUCell score either >1.2 (pathway expression increased) or <0.8 (pathway expression decreased). From the enriched metabolomics pathways, we found that 31 pathways (22.1%) increased, and 76 pathways (54.3%) decreased gene expression. From the selected lipidomics KEGG and BioCyc terms, we found that 8 pathways (61.5%) increased, and 1 pathway (7.7%) decreased gene expression. Out of all lipid reactions, we found 117 reactions (75%) increased, and 13 reactions (8.3%) decreased gene expression.

We next identified for each increased and decreased metabolomic and lipidomic pathway all up- and downregulated DEGs between Xbp1s and control flight muscle. Many DEGs were shared between annotated pathways and reactions, which was expected as many KEGG and BioCyc terms we selected either overlapped or were redundant, as was true for many of the 117 LipidSig reactions. We next clustered these DEGs into three broad categories: 1) upregulated lipid metabolism (43 upregulated DEGs; **Supplemental Figure 13A**), 2) upregulated non-lipid metabolism (39 upregulated DEGs; **Supplemental Figure 13B**), and 3) downregulated non-lipid metabolism (90 downregulated DEGs; **Supplemental Figure 13C**). We did not include a downregulated lipid metabolism category as most of the lipid-associated transcriptional changes were upregulated. We next subcategorized these broad pathways into more specific metabolic pathways and allowed DEGs to be present in as many pathways as annotated: 1) upregulated lipid metabolism: glycerophospholipid metabolism (21 upregulated DEGs) and other lipid metabolism (mostly sphingolipid and GPI-anchor metabolism; 28 upregulated DEGs); 2) upregulated non-lipid metabolism: glutathione and methionine metabolism (15 upregulated DEGs), pyrimidine metabolism (12 upregulated DEGs), and other non-lipid metabolism (14 upregulated DEGs); 3) downregulated non-lipid metabolism: carbohydrate metabolism (45 downregulated DEGs), arginine and proline metabolism (12 downregulated DEGs), purine metabolism (16 downregulated DEGs), and retinol metabolism (15 downregulated DEGs).

Finally, we calculated the percentage of DEGs from each broad or specific metabolic category that were found in SCENIC-defined differentially active TF regulons. Overall, we found substantial enrichment of many DEGs within regulons, and that DEGs of upregulated metabolic pathways were almost entirely specific to TFs with increased activity, while DEGs of downregulated pathways were specific to TFs with decreased activity (**Figure 5B**). Xbp1, cnc, Atf6, Cf2, and Mitf were most enriched for upregulated lipid and non-lipid DEGs, while FoxO, FoxK, and REPTOR-BP were most enriched for downregulated non-lipid DEGs. SREBP and CrebB were specifically enriched for glycerophospholipid DEGs. We also determined the percent DEG overlap with SCENIC regulons built using only Xbp1 activated flight muscle cells (**Supplemental Figure 10B**). Similar results were found for Xbp1, cnc, CrebB, Cf2, and FoxO, with FoxO including several upregulated metabolic pathways as well (**Supplemental Figure 14A**).

We next wanted to validate our findings using a TF regulon construction tool other than SCENIC. We chose high dimensional weighted gene co-expression network analysis (hdWGCNA), a framework similar to SCENIC that uses co-expression network analysis to build gene modules assigned to TFs with high binding predictions (Morabito et al. 2023). Unlike our differentially active SCENIC regulons, our hdWGCNA regulons had near equal representation of positively regulated and negatively regulated target genes for each TF (i.e., genes predicted to be targeted by a specific TF that increased in co-expression (positively regulated) or decreased (negatively regulated)). We divided each regulon into positively and negatively regulated predicted targets, then selected regulons shared with our SCENIC analysis, and determined percent metabolic DEG overlap for each TF (**Figure 5C**). Similar to SCENIC, hdWGCNA predicted that Xbp1, SREBP, cnc, crc, and Cf2 positively regulated DEGs of upregulated metabolic networks, while FoxO and doublesex (dsx) positively regulated DEGs of downregulated metabolic networks (**Figure 5C**).

Interestingly, SREBP and cnc were predicted to negatively regulate downregulated metabolic DEGs, while FoxO, dsx, and BTB-protein-VII (BtbVII) were predicted to negatively regulate upregulated metabolic DEGs, suggesting that some GRNs may antagonize each other by positively or negatively regulating each other’s target metabolic networks. This was supported by determining DEG overlap with specific metabolic pathways, in which we found that SREBP and cnc were predicted to negatively regulate carbohydrate metabolism while FoxO and dsx were predicted to negatively regulate lipid metabolism (**Supplemental Figure 14B**). Altogether, across three strategies to predict GRN activities (two SCENIC approaches and hdWGCNA-defined positive and negative targets), we concluded that increased Xbp1, SREBP, cnc, and Cf2 activity likely account for upregulated lipid and nonlipid metabolism, while decreased FoxO activity likely accounts for downregulated nonlipid metabolism, namely carbohydrate pathways.

To interrogate the relationships between these GRNs, we further analyzed hdWGCNA-defined negatively regulated FoxO targets and found significant KEGG term enrichment of beta-oxidation and acyl-CoA synthesis, fatty acid biosynthesis and degradation, and inositol phosphate metabolism (**Supplemental Figure 15A**). hdWGCNA-defined negatively regulated SREBP and cnc targets were enriched for KEGG terms related to glycolysis, citrate/TCA cycle, pentose phosphate pathway, and oxidative phosphorylation (**Supplemental Figure 15B, 15C**). In summary, our assignment of metabolomic and lipidomic changes to GRNs predicted that a handful of TFs downstream of ER stress (Xbp1, SREBP, cnc, crc, Cf2) control upregulated metabolic networks and work to suppress downregulated networks, while decreased FoxO activity (and potentially others like FoxK, REPTOR-BP, dsx) may contribute to downregulated metabolic pathways.

### Chronic SREBP activation degrades muscle and disrupts carbohydrate metabolism

Throughout our analyses, SREBP was consistently predicted to be the most active metabolic regulator alongside Xbp1 in muscle. While the consequences of SREBP activation have been extensively studied in hepatocytes and adipose tissue, much less is characterized about its function in skeletal muscle (Shimano et al. 2017). In flies, aberrant SREBP activation and dysregulation of phospholipid metabolism in cardiac muscle was shown to increase heart failure (Lim et al. 2011). As we observed both SREBP and glycerophospholipid metabolism increased downstream of ER stress, we hypothesized that SREBP activation alone in muscle would be detrimental for the tissue and animal. To test this, we expressed a constitutively active mutant of SREBP under UAS (UAS-SREBP.Cdel, hereby referred to as “SREBP[ACT]”) in which the sequence is truncated before the first transmembrane domain, preventing its ER membrane retention, and results in constitutive localization to the nucleus.

Using a temperature sensitive pan-muscle driver, we observed a sharp decline in lifespan after persistent SREBP[ACT] expression (**Figure 6A**). Flies also rapidly lost their ability to climb in response to negative geotaxis assays in the first few days of SREBP activation (**Figure 6B**). Moreover, we found that the vast majority of myofibers in thorax muscles were degraded after immunohistochemical staining for actin organization (**Figure 6C**). These results indicated that overactivation of SREBP is highly damaging to muscle and organismal health. Our metabolomic, lipidomic, and transcriptomic analyses predicted that SREBP not only activates lipogenic programs in muscle, but that it may interfere with carbohydrate metabolism (**Supplemental Figure 15B**). Consistent with this, we found significantly increased whole-body glucose and glycogen levels after 3 days of SREBP muscle activation (**Figure 6D**). We also observed decreased protein levels, further supporting myofiber degradation and catabolism.

**Figure 6.**
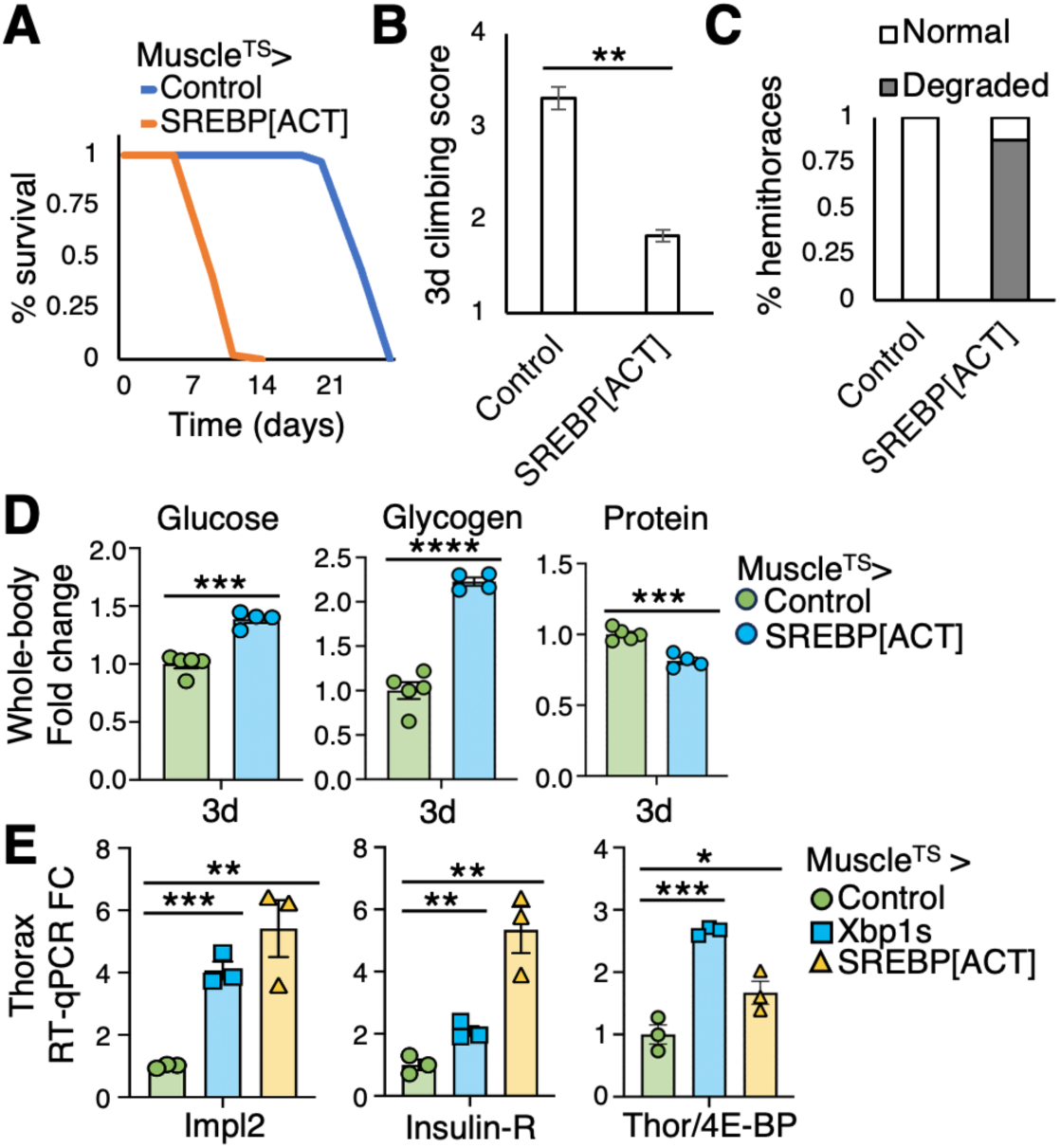
Constitutive SREBP activation in muscle quickly diminishes muscle function and lifespan, while disrupting glucose metabolism and insulin signaling. **A)** Adult male lifespan monitoring with sustained UAS-SREBP[ACT] (constitutively active SREBP mutant) expression in muscle. **B)** Negative geotaxis climbing score (ranging 1-4) after 3d SREBP activation. **C)** Percentage of dissected hemithoraces with degraded muscle myofibers after 3d SREBP activation, determined by phalloidin actin staining patterns and scored with confocal microscopy. **D)** Whole-body detection of glycogen, glucose, and protein after 3d SREBP activation; 1 rep = 8 flies. **E)** RT-qPCR mRNA expression fold change of Impl2, Insulin receptor (Insulin-R), and Thor/4E-BP in thoraces after 4d SREBP or Xbp1 activation; 1 rep = 10 thoraces. All significance determined using unpaired T-test.

Mammalian SREBP is known to be regulated by insulin signaling and influence glucose metabolism, including muscle cells (Nadeau et al. 2004, DeBose-Boyd et al. 2019). Flies however lack obvious Insulin-induced genes (INSIGs) that mediate mammalian insulin regulation of SREBP activation, and the relationship between insulin signaling and SREBP in *Drosophila* is less clear (Dobrosotskaya et al 2003, DiAngelo et al. 2009). The increased glucose and glycogen after SREBP[ACT] expression prompted us to measure the expression of insulin signaling components in muscle. We performed RT-qPCR on dissected thoraces after 3 days of SREBP activation and found significantly increased expression of insulin receptor (InR), Thor/4E-BP, and Impl2, an insulin-like peptide (ILP) binding protein that prevents InR recognition and signaling, suggesting disrupted and decreased insulin signaling in muscle (**Figure 6E**). We also observed InR, Impl2, and Thor/4E-BP upregulation in Xbp1 activated muscle. Overall, these data suggest that aberrant SREBP activity in muscle degrades tissue and disrupts insulin signaling and carbohydrate metabolism.

## Discussion

Our data and analyses revealed that Xbp1 splicing and activation during protein misfolding in the ER can substantially rewire muscle metabolism through various potential transcriptional networks (**Figure 7A**). Foremost, we found that Xbp1 itself likely binds hundreds of targets that promote lipogenesis and fatty acid utilization, induce antioxidant programs, and affect myofiber actin cytoskeletal dynamics and circadian rhythm (**Supplemental Figure 4A, 4B, Supplemental Figure 10C)**. GRN analyses revealed that other TFs, such as SREBP, cnc, crc, Cf2, and CrebB, may co-regulate these changes. We found that SREBP, known as a master lipogenesis regulator, alone can degrade muscle and disrupt carbohydrate metabolism. Along with cnc, SREBP activity downstream of ER stress may inhibit expression of core carbohydrate and oxidative phosphorylation genes (**Supplemental Figure 15B, 15C**). Across all our analyses, we also predicted decreased FoxO activity, which likely further contributes to downregulating sugar and starch metabolism and mitochondrial respiration. FoxO was also predicted to actively downregulate lipogenesis and fatty acid beta-oxidation genes (**Supplemental Figure 15A**). These data suggested an interesting antagonism between TFs that control lipid metabolism (e.g., Xbp1, SREBP) and those that control carbohydrate metabolism (e.g., FoxO) which is exploited during times of cellular stress to alter energy demands and utilization, potentially impacting the overall health of muscle (**Figure 7A**).

**Figure 7.**
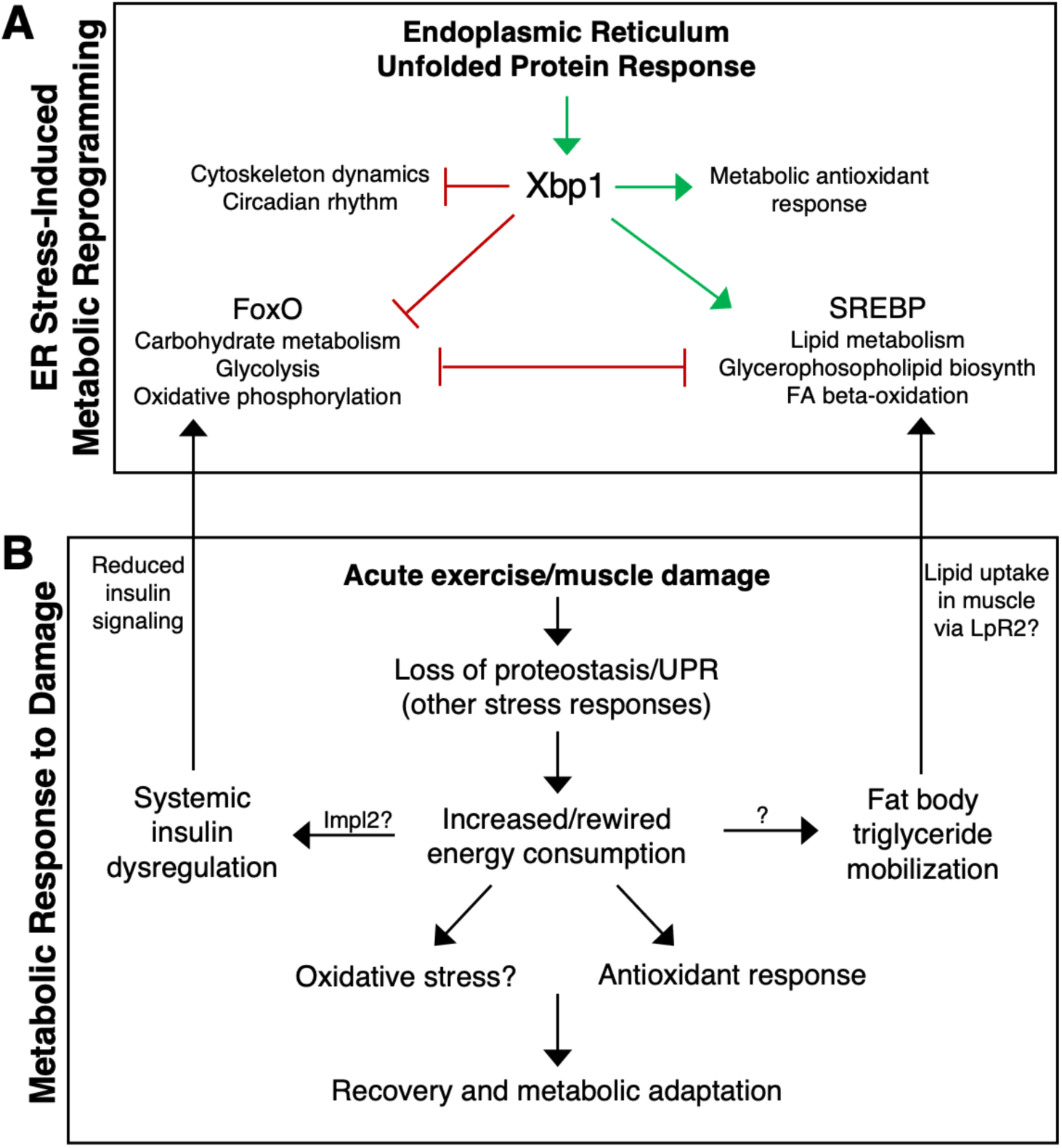
Proposed models of ER stress-induced transcriptional reprogramming of muscle metabolic networks. **A)** Xbp1 activation downregulates genes associated with cytoskeleton dynamics and circadian rhythm while upregulating a multifaceted antioxidant metabolic response. Xbp1 also upregulates GRNs, likely along with co-regulators such as SREBP, that control lipid metabolism and glycerophospholipid biosynthesis. FoxO activity decreases, along with genes and metabolic processes related to glycolysis, glucose metabolism, and oxidative phosphorylation. GRNs that transcriptionally regulate carbohydrates/oxidative phosphorylation and lipids/fatty acid oxidation may antagonize each other to influence the overall metabolic wiring of muscle tissue. **B)** In the context of muscle damage, such as acute exercise, proteostatic stress responses transcriptionally alter energy consumption and fuel sources while simultaneously affecting systemic insulin signaling and adipose tissue triglyceride storage and mobilization. Consequences of metabolic rewiring and fuel switching may be increased oxidative stress along with a robust antioxidant response. Ultimately, these transcriptionally regulated metabolic networks may allow for metabolic adaption in muscle after repair and recovery. This could be beneficial, as with endurance exercise training, or detrimental, as with maladaptation and increased susceptibility to metabolic disease.

FoxO proteins are widely accepted pro-longevity factors that promote healthy aging, insulin signaling, and autophagy, from worms to mammals (Martins et al. 2016; Davy et al. 2018). However, its role in skeletal muscle health and disease has been more complex. Several studies, including our own, have shown the potential benefits of FoxO activity in regulating muscle metabolism, function, and disease susceptibility (Demontis et al. 2020; Bai et al. 2013; Saavedra et al. 2023; Sanchez et al. 2014). Conversely, its catabolic and autophagic functions can exacerbate muscle atrophy and wasting disorders (Sandri et al. 2004; Chen et al. 2022; Judge et al. 2014). These diverse outcomes have made FoxO proteins desirable therapeutic targets for various tissues and diseases (Orea-Soufi et al. 2022). As with most stress-responsive factors, FoxO activity most likely needs to be modulated and balanced in muscle to limit aging and disease, whereas overactivation is detrimental to the tissue. Our data suggest that FoxO activity decreases during muscle ER stress and contributes to disrupted glycogen and glucose homeostasis. This is supported by studies suggesting Xbp1 directly suppresses FoxO in several tissues and species (Zhao et al. 2021; Kishino et al. 2017; Schiattarella et al. 2021). We previously reported that increased FoxO in muscle can prevent age-associated loss of proteostasis and improve healthspan (Demontis et al. 2010). Further experimentation will be necessary to confirm if increasing FoxO activity during ER stress can promote muscle health and metabolism, especially in the context of aberrant phospholipogenesis and SREBP activation.

Increased lipogenesis, namely phospholipids, has been shown to be an essential part of UPR and Xbp1 function to expand ER membrane to ease the burden of abundant protein secretion (Fagone et al. 2009). The physiological consequences of sustained phospholipid biosynthesis in skeletal muscle have not been fully explored. In fly heart tissue, SREBP activation and phospholipid metabolism disruption led to cardiac dysfunction (Lim et al. 2011). The possible physiological benefits of Xbp1/SREBP phospholipogenesis in muscle may include adapting the tissue for increased secretory capacity, as skeletal muscle acts as an endocrine organ secreting many myokines in response to exercise (Hoffman et al. 2017). This endocrine function is critical for muscle and organismal adaptation to exercise. Indeed, we observed altered insulin signaling in muscle and TG degradation in adipose tissue after muscle Xbp1 activation, suggesting interorgan crosstalk. Released free fatty acids from adipose tissue may act as phospholipid building blocks and beta-oxidation fuel sources in muscle during times of increased energy demand and damage (e.g., exercise). If these transcriptional programs are sustained, as what we report here, metabolic rewiring from glucose to lipid metabolism may ultimately degrade muscle, as we showed with constitutive SREBP activation (**Figure 6B, 6C**). We recently reported that prolonged activation of the transcription factor Repressed by TOR (REPTOR) in the context of tumor-induced muscle wasting decreases glycolysis and degrades muscle fibers via increased utilization of fatty acid oxidation (Saavedra et al. 2023). Therefore, the transcriptional reprogramming of muscle metabolic plasticity downstream of cellular stress is a complex and critical mechanism to elucidate.

More broadly, we propose that some kinds of muscle cellular damage, such as after acute exercise, cause a proteostatic and ER stress response (and likely other responses, such as inflammation) that increases and alters energy consumption, namely utilization of glucose vs fatty acids (**Figure 7B**). This could lead to increased oxidative stress and antioxidant responses, changes in insulin signaling (potentially through secretion of Impl2), and mobilization of triglyceride storage in adipose tissue. Decreased insulin signaling and increased uptake of circulating free fatty acids in muscle (via upregulation of LpR2), may contribute to the regulation and activity of GRNs controlling carbohydrate vs lipid metabolism (**Figure 7B**). Ultimately, muscle cells must recover from damage and return to homeostasis. The utilization of GRNs in our findings suggest that metabolic rewiring in response to damage is largely transcriptionally controlled. We thus favor a model in which long-term metabolic adaptation to muscle damage, such as regular exercise training, are in part driven by stress responsive GRNs.

## Materials and Methods

### Drosophila stocks and husbandry

Flies were maintained on standard lab food with no more than 20 males or females per vial. Crosses using temperature-sensitive GAL80 control of GAL4 were maintained at 18°C through development under a 12hr/12hr light/dark cycle. After eclosion, adult flies were maintained at 18°C for 2-3 days for mating and maturity (all female flies used in experiments were mated). Flies were transferred to 29°C for experiments to permit GAL4 activity and UAS-transgene expression and flipped onto fresh food every 1-2 days.

Unless otherwise noted, all experiments with Muscle^TS^ driver were performed with Perrimon lab stock of *tubulin-Gal80ts; Mhc-Gal4*. Other muscle drivers included lab stocks of *tubulin-Gal80ts; dMef2-Gal4*, and *tubulin-Gal80ts; Act88F-Gal4*. Stocks crossed as control flies for single-nuclei RNA-sequencing: *OregonR* (Perrimon lab stock). Stocks crossed as control flies for all other experiments: *UAS-emptyVK37* (gift from Hugo Bellen). Other UAS stocks: *UAS-Xbp1-RB* (“UAS-Xbp1s”, gift from Hyung Don Ryoo), *UAS-Rh1^G69D^* (gift from Hyung Don Ryoo), *UAS-SREBP.Cdel* (Bloomington stock # 8244).

### Climbing score, lifespan, starvation, heat shock

Climbing scores were determined using negative geotaxis assays. Flies were transferred to two new vials taped end-to-end with halfway marks noted on each vial, segmenting the climbing chamber into four quadrants. Flies were tapped down and imaged after 10 sec. Number of flies in each quadrant were counted and scored (lowest quadrant = climbing score of 1; highest quadrant = climbing score of 4). This was repeated three times and scores were averaged across triplicate runs. 3-4 biological replicates (each containing 20 flies) were recorded per genotype. Final climbing score was calculated for each biological replicate by multiplying percentage of total flies with each score by the score itself and summing: ((% flies scored 1) x 1) + ((% flies scored 2) x 2) + ((% flies scored 3) x 3) + ((% flies scored 4) x 4) = Final Climbing Score. Final scores ranged from 1 (100% of flies in the lowest quadrant) to 4 (100% of flies in the highest quadrant).

For lifespan analyses, survival was calculated by the percentage of flies still alive at a given time point over the starting number. Flies were flipped onto new food every 1-2 days. Dead flies remaining in old vials were recorded as dead. Wet starvation assays were performed by transferring flies from standard food to vials containing only 1% agar. Fly death was recorded regularly until all flies perished. Heat shock assays were performed by transferring flies (20/vial) into fresh vials with no food and fully submerged in a 35°C water bath for 30 min to prime heat shock responses. They were then immediately transferred to a 39°C water bath and recorded for “paralysis”/inactivity every 4 min until all flies were inactive.

### Triglyceride, glycogen, glucose, and protein measurements

Whole flies with heads removed (to avoid eye pigment influencing the assays) were used for colorimetric metabolite assays. For males, 8 flies were used per replicate; for females, 6 flies were used per replicate. Flies were collected in 1.5 mL tubes, flash frozen in liquid nitrogen, and stored at -80°C until processing. Flies were homogenized in 150 uL 0.1% Triton X-100/1X PBS using 1 mm zirconium oxide beads (Next Advance Lab Products, ZROB10) in a TissueLyser II homogenizer (Qiagen). Homogenate was collected and centrifuge at 3,000g for 5 min. 2 uL was used for protein detection with Pierce BCA Protein Assay Kit (Thermo Fisher Scientific, 23227) and incubated at 37°C for 30 min before reading with a spectrophotometer. 10 uL was used for glycogen and glucose measurement. Glycogen measurements were first incubated at 37°C for 30 min with 1/100 amyloglucosidase (Sigma-Aldrich, A1602). Glucose and glycogen (after amylogucosidase treatment) were detected using Infinity Glucose Hexokindase Reagent (Thermo Fisher Scientific, TR15421). 5 uL of supernatant was used for triglyceride measurement using Triglycerides Reagent (Thermo Fisher Scientific, TR22421).

### mRNA extraction and reverse transcription qPCR

10 thoraces per biological replicate were dissected in 0.3% Triton X-100/1X PBS and lysed in 200 uL TRIzol (Thermo Fisher Scientific) with 1 mm zirconium oxide beads (Next Advance Lab Products, ZROB10) in a TissueLyser II homogenizer (Qiagen). Homogenized samples were stored in TRIzol at -80°C until extraction. mRNA was extracted after bringing TRIzol volume to 1 mL total and per TRIzol manufacturer’s protocol. Extracted RNA was measured using NanoDrop and equal amounts across samples were added to reverse transcription synthesis of complementary DNA using iScript cDNA Synthesis Kit (Bio-Rad, 1708890) according to manufacturer’s protocol. qPCR was performed using a Thermal Cycler CFX 96 Real-Time System qPCR machine and amplified with iQ SYBR Green Supermix (Bio-Rad). Primers used were as follows. alpha-Tubulin (housekeeping gene for normalization): FWD CAACCAGATGGTCAAGTGCG, REV ACGTCCTTGGGCACAACATC; spliced Xbp1 (Xbp1s): FWD TGGATCTGCCGCAGGGTAT, REV GCGCTTGACGTCGAACTCTT; Hsc70-3: FWD GATTTGGGCACCACGTATTCC, REV GGAGTGATGCGGTTACCCTG; Impl2: FWD AAGAGCCGTGGACCTGGTA, REV TTGGTGAACTTGAGCCAGTCG; Insulin receptor (InR): FWD AAGCGTGGGAAAATTAAGATGGA, REV GGCTGTCAACTGCTTCTACTG; 4E-BP/Thor: FWD TCCTGGAGGCACCAAACTTATC, REV GGAGCCACGGAGATTCTTCA.

### Immunohistochemistry

Cuticle fat body was dissected after 10d Xbp1s muscle expression and fixed in 4% paraformaldehyde and permeabilized in 0.1% Triton X-100/1X PBS for 30 min and washed three times to thoroughly remove Triton X-100. 1 mg/mL stock of BODIPY-488 in DMSO was diluted 1/500 in PBS and stained cuticles for 30 min at room temp. Tissues were washed thoroughly in PBS and mounted on slides in VectaShield with DAPI (1200).

For myofiber degradation, dissected thoraces were fixed in 4% paraformaldehyde for 1 hr, bisected with a razor blade, and stained for 2 hr at room temp with Phalloidin-AF555 (1:200, Thermo Fisher Scientific A3405%). Hemithoraces were then mounted on u-Dish 35 mm high Glass Bottom (Ibidi) in VectaShield. Z stack acquisition of thoraces on the Yokogawa CSU-W1 Spinning disk with 20x objective. Myofiber degradation was scored as previously described (Saavedra et al. 2023) using actin/Phalloidin staining pattern to interpret sarcomere structure.

### Whole-body single-nuclei RNA-sequencing prep

Whole-body single-nuclei RNA-sequencing (snRNA-seq) was performed as previously described (Liu et al. 2025). Briefly, 16 male flies (heads removed) after 3d of Xbp1s expression at 29°C were flash frozen in liquid nitrogen and stored at -80°C for two weeks until processing. Frozen flies were transferred to 1 mL Dounce in lysis buffer (250 mM sucrose, 10 mM Tris pH 8.0, 25 mM KCl, 5 mM MgCl, 0.1% Triton-X, 0.5% RNasin plus (Promega, N2615), 1× protease inhibitor (Promega, G652A), 0.1 mM DTT) and homogenized and filtered as previously described. After centrifugation and washing, nuclei were filtered again immediately before FACS sorting. Nuclei were stained with DRAQ7TM Dye (Invitrogen, D15106) and sorted using Sony SH800Z Cell Sorter. Nuclei were then collected and resuspended at 700–800 cells/uL in 1X PBS with 0.5% BSA and 0.5% RNasin plus. Immediately after, 10X Genomics protocol (Chromium Next GEM Single Cell 3’_v3.1_Rev_D) was followed for single-cell library preparation. Two reactions per genotype (20,000 nuclei total) were processed and sequenced using Illumina NovaSeq 6000 S1.

### snRNA-seq Data Analysis

Raw sequencing data were aligned to the *Drosophila melanogaster* reference genome (BDGP6.32, Ensembl release 104) using 10x Genomics Cell Ranger v7.1.0. Ambient RNA contamination in the resulting count matrices was estimated and corrected using SoupX (v1.6.2) (Young et al. 2020) with default parameters in R (v4.5.0). The corrected gene-by-cell expression matrices were processed in Seurat (v5.2.1). Nuclei with fewer than 500 unique molecular identifiers (UMIs), 300 genes and >10% mitochondrial content were excluded as low quality. Suspected multiplets were filtered based on data-driven thresholds: nuclei were removed if their nUMI or nGene counts exceeded the median plus five times the median absolute deviation of the dataset. Genes which were not expressed in at least 10 cells across the dataset were filtered out.

The data was then normalized using NormalizeData and a total of 2,000 highly variable genes were identified using FindVariableFeatures. The data was scaled, and dimensionality reduction was performed by principal component analysis (50 principal components). Clustering was performed using Seurat’s FindNeighbors and FindClusters functions (resolution 0.4), and Uniform Manifold Approximation and Projection (UMAP) was applied to visualize the data. Clusters were annotated based on canonical marker gene expression, identified with Seurat’s FindAllMarkers function.

Differential gene expression between Xbp1s and control cells was performed within each cell type using the Wilcoxon rank-sum test. Genes with absolute log2 fold change > 1 and p-value < 0.05 were considered differentially expressed. Overrepresentation analysis for pathway enrichment was performed using PAthway, Network, and Gene-set Enrichment Analysis (PANGEA) (Hu et al. 2023). Pathway databases used for enrichment tests included Gene Ontology: Biological Process (GO:BP), Kyoto Encyclopedia of Genes and Genomes (KEGG), and BioCyc.

### SCENIC and hdWGCNA regulon construction

For each cell type and condition, gene expression count matrices and cell metadata were generated. Due to the stochastic nature of GRN inferencing step, SCENIC (v1.3.1) was run 20 times per cell type–condition group, each tune with a different random seed. For each run, SCENIC performed gene regulatory network inference, regulon prediction, and activity scoring. Regulons were retained if detected in at least 80% of the runs for a group. Similarly, for each regulon, only target genes present in at least 80% of the runs for that regulon were kept. This resulted in a consensus list of regulons and their target genes for each tissue-condition combination, which was used in subsequent analyses.

Co-expressed gene modules were identified using hdWGCNA (v0.4.05) separately for each tissue–condition group. Briefly, for each tissue–condition expression matrix, metacell objects were constructed (k = 25, max_shared = 10, min_cells = 75, target_metacells = 1000) and modules were detected according to hdWGCNA’s default workflow. Promoter regions (2 kb upstream of transcription start sites) of protein-coding genes were scanned for transcription factor binding motifs using the JASPAR 2024 Drosophila melanogaster motif database. To infer regulatory networks, ConstructTFNetwork() function was run 20 times per group and regulons were retained if detected in at least 80% of runs. For each regulon retained, target genes present in at least 80% of the runs for that regulon were kept.

### AUCell scoring

AUCell (v1.3.0) was applied to tissue-specific gene expression matrices to assess regulon activity. AUCell was run using an aucMaxRank parameter corresponding to the top 5% of expressed genes per cell. For each regulon or pathway term, AUCell scores were averaged across cells within the Xbp1s and control clusters for downstream comparisons.

### Targeted metabolomics and untargeted lipidomics prep

Metabolomics and lipidomics were performed on the same thoraces after 6d of either Xbp1s or Rh1^G69D^ muscle expression. 25 thoraces per biological replicate were dissected in 1X PBS, flash frozen in liquid nitrogen, and stored at -80°C until processing. To extract polar and nonpolar metabolites for targeted metabolomics and untargeted lipidomics, respectively, frozen thoraces were transferred to 1 mL Dounce homogenizer (TIGHT) with 600 uL chloroform and 300 uL methanol (2:1 chloroform-methanol final). Samples were homogenized on ice in the Dounce until fully dissociated and transferred with a glass pipet to a 15 mL glass tube with Teflon cap (no plastic pipets or tubes were used throughout the protocol). An additional 600 uL chloroform and 300 uL methanol was added to final volume of 1.8 mL. Samples were mixed by vortexing and rotated at room temperature for 30 min. 360 uL (0.2 volumes) of HPLC-grade deionized water was added and vortex three times. Centrifugation at 1000 x g, 4°C for 10 min separated upper aqueous phase (polar metabolites) and lower phase (nonpolar lipid metabolites). Upper phase was slowly collected using a glass pipet and transferred to microcentrifuge tubes to evaporate overnight in a SpeedVac vacuum rotary evaporator. Dried samples were stored at -80°C until submission for mass spec. Lower phase was transferred using a glass pipet to a 1.85 mL glass vial and dried under nitrogen gas to prevent lipid oxidation. They were then stored at -80°C under sumitting for mass spec. Polar metabolomics were resuspended in 20 uL HPLC-grade water and injected and analyzed using a hybrid 6500 QTRAP triple quadrupole mass spectrometer (AB/SCIEX) connected to a Prominence UFLC HPLC system (Shimadzu). Nonpolar lipidomics were resuspended in 35 uL HLPC-grade 50% methanol/50% isopropyl alcohol (IPA) and injected and analyzed using untargeted high-resolution LC-MS/MS on a Thermo QExactive Plus Orbitrap mass spectrometer.

### Metabolomics and Lipidomics Data Analysis

For targeted metabolomics, integrated peak intensities were obtained from raw mass spectrometry data and imported to R. Metabolites detected in both positive and negative ionization modes were de-duplicated by retaining the ion mode with the higher median peak intensity in wild-type samples. Metabolites missing in more than 20% of samples in the control, Xbp1s, and Rh1^G69D^ groups were excluded from further analysis. Remaining missing values were imputed by replacing them with half the minimum non-missing value observed for that metabolite across all samples. For downstream statistical and pathway analysis, data were imported into MetaboAnalyst 6.0 and processed using sample median normalization and log10 transformation.

For untargeted lipidomics, integrated peak intensities were obtained from raw mass spectrometry data as previously described and imported to R. Lipids detected with multiple adducts were de-duplicated by retaining the adduct species with the highest median peak intensity in control thoraces. Lipid names were standardized using LipidSig 2.0, which was also used for subsequent downstream statistical analysis.

## Acknowledgements

We thank Dr. John M. Asara at the Beth Israel Deaconess Medical Center for his help performing mass spectrometry metabolomics and lipidomics and subsequent data analysis. We thank Dr. Hyung Don Ryoo for gifting UPR-inducing transgenic flies and providing critical project feedback. We thank Dr. Richard Binari and all members of the Perrimon lab for invaluable research support and assistance.

## Author Contributions

JP and NP conceived this study. JP designed and conducted the majority of experiments. SS conducted SREBP activation experiments. MQ performed all bioinformatic analyses. MQ, JP, and YH developed bioinformatic pipelines. JP wrote this manuscript. MQ, NP, and YH edited this manuscript.

## Notes

### Competing Interest Statement

The authors have declared no competing interest.

