## Supplemental Figures for "Protein folding stress transcriptionally reprograms muscle metabolism"

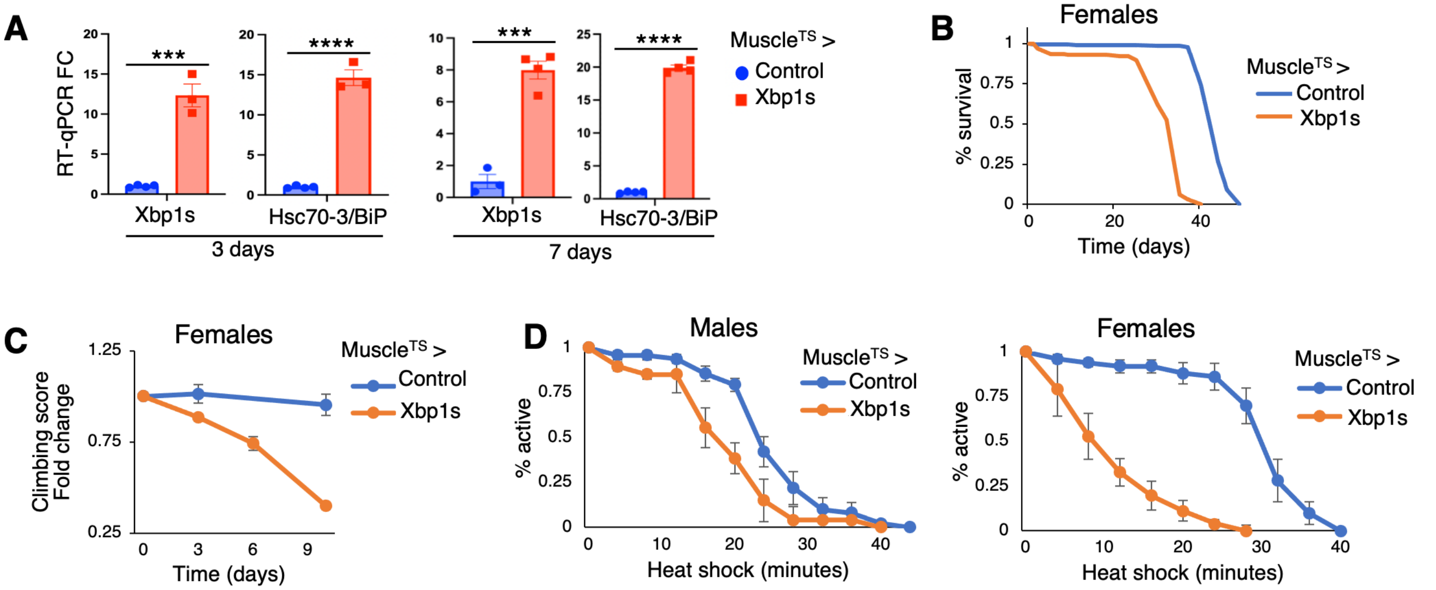


**Supplemental Figure 1. Characterizing organismal consequences of muscle-specific Xbp1 activation in males and females. A)** Thorax RT-qPCR mRNA expression fold change of spliced Xbp1 (Xbp1s) and downstream target gene Hsc70-3/BiP after 3d and 7d expression of Xbp1s in adult muscle; 1 rep = 10 thoraces. **B)** Adult female lifespan monitoring with sustained Xbp1 activation in muscle. **C)** Negative geotaxis climbing score (ranging 1-4) fold change (over 0d time point) in adult females over time. **D)** Heat shock sensitivity assay after 6d Xbp1s muscle expression in males (left) and females (right). Vials containing flies were transferred to 39°C water baths and number of inactive flies were recorded every 4 min. All significance was determined using unpaired T-tests.

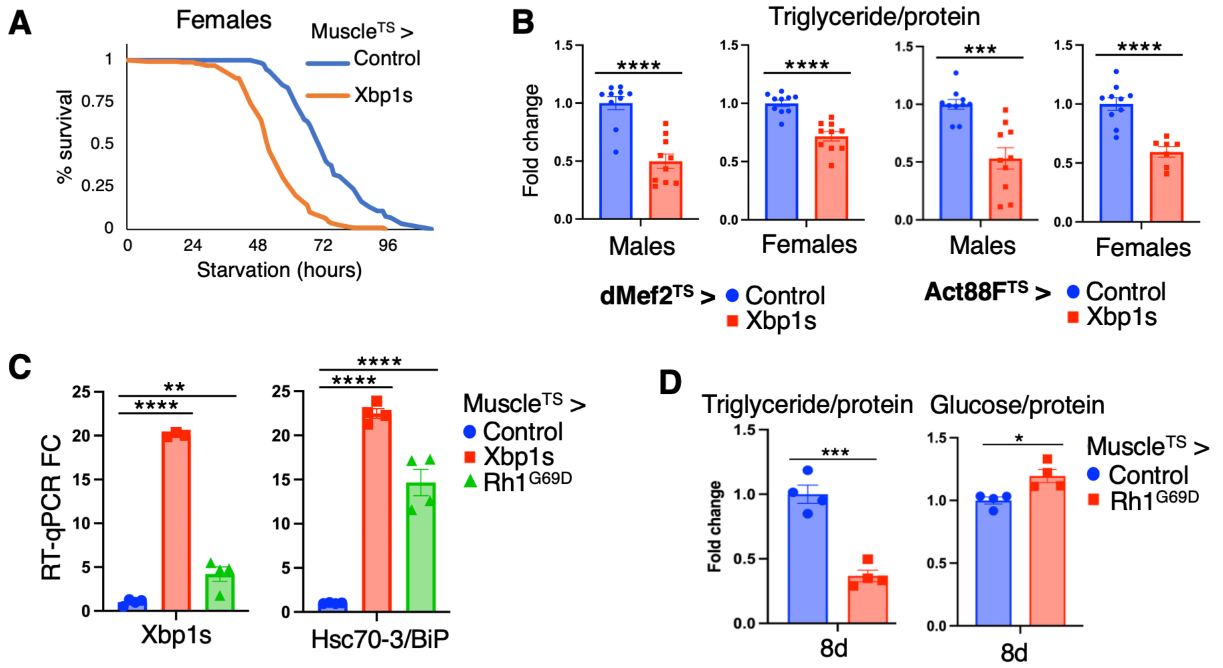


**Supplemental Figure 2. Muscle-specific UPR induction causes whole-animal depletion of macronutrient storage. A)** Wet starvation (1% agar) survival of adult females after 6d Xbp1s muscle expression. **B)** Whole-body measurement of triglyceride content, normalized to total protein, in adult male and female flies after 9d Xbp1s expression in muscle. Two distinct, temperature-sensitive (TS) muscle GAL4 drivers were used, dMef2 and Act88F. For males, 1 rep = 8 flies; for females, 1 rep = 6 flies. **C)** Thorax RT-qPCR mRNA expression fold change of spliced Xbp1 (Xbp1s) and downstream target gene Hsc70-3/BiP after 3d of Xbp1s or misfolded rhodopsin (Rh1$^{G69D}$) muscle expression; 1 rep = 10 thoraces. **D)** Whole-body measurement of triglyceride and glucose content, normalized to total protein, after 8d Rh1$^{G69D}$ muscle expression.

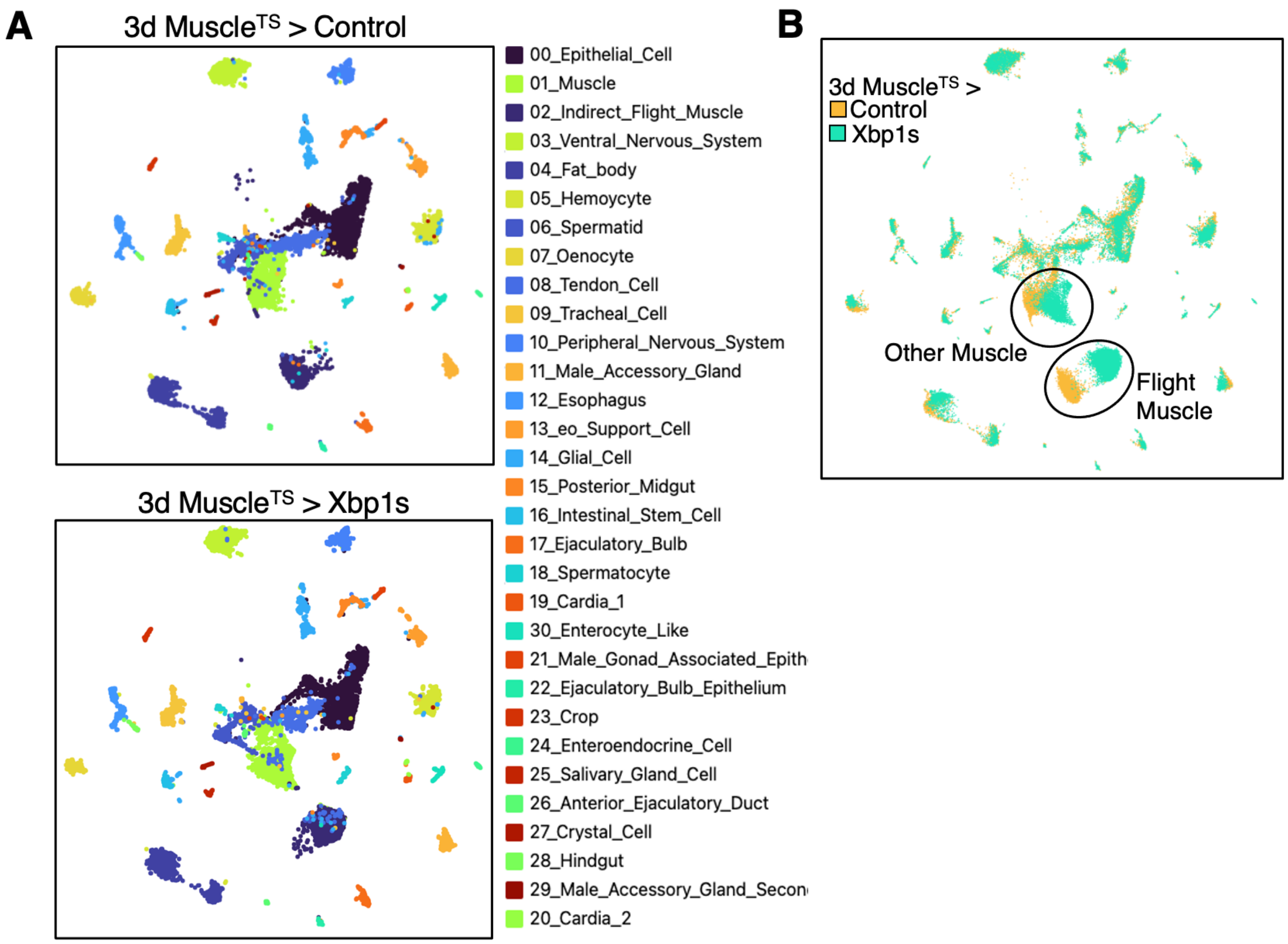


**Supplemental Figure 3. Characterizing whole-body single-nuclei RNA-sequencing (snRNA-seq) cell clusters. A)** UMAP of whole-body (heads removed) snRNA-seq from adult male flies after 3d Xbp1s muscle expression. 31 distinct clusters were identified and annotated based on published tissue marker genes. Equal tissue representation and cell numbers were found between control and Xbp1 activated samples. **B)** UMAP overlaying snRNA-seq clusters from control and Xbp1s flies. Generally, significant overlap was observed for all clusters except flight muscle and other muscle, in which Xbp1s-expressing cells notably shifted from control cells. This indicates impactful transcriptomic changes between the two conditions for both muscle clusters.

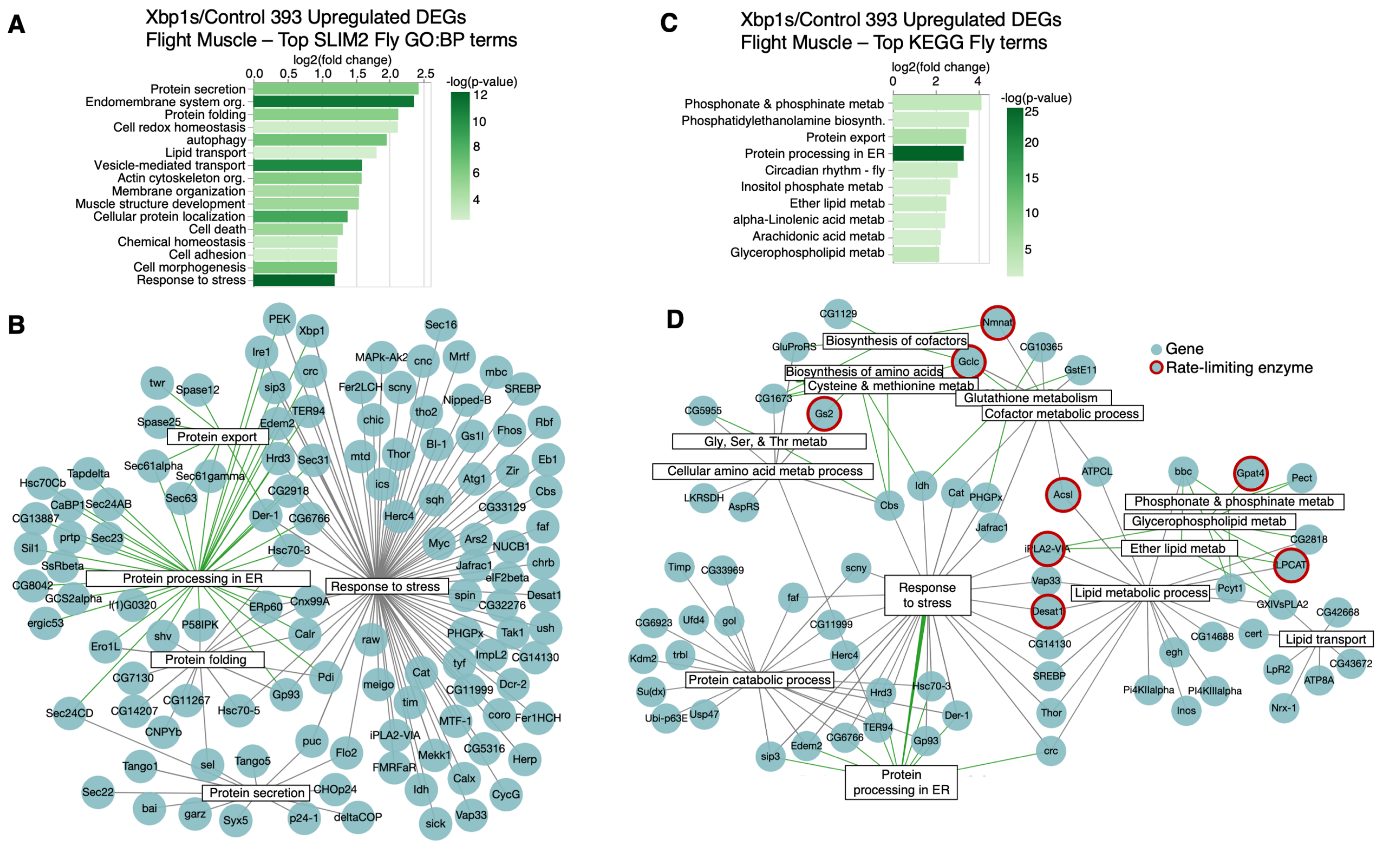


**Supplemental Figure 4. Gene Ontology (GO) and KEGG enrichment of upregulated differentially expressed genes (DEGs) in Xbp1-activated flight muscle. A)** PAthway, Network, and Gene-set Enrichment Analysis (PANGEA) overrepresentation analysis of *Drosophila* SLIM2 GO Biological Process (BP) terms for all upregulated DEGs (393 genes total) in flight muscle. Top GO BP term enrichment shown. **B)** Gene node plots for select enriched terms in Xbp1s flight muscle upregulated DEGs. Each gene/node represents a significantly upregulated DEG. Interconnectedness between ER stress terms, “protein export”, “protein processing in ER”, “protein folding”, and “protein secretion”, and general “response to stress” is visualized. **C)** Overrepresentation analysis showing most enriched *Drosophila* KEGG terms in upregulated flight muscle DEGs. **D)** Gene node plots for select enriched terms in Xbp1s flight muscle upregulated DEGs, showing connectedness between metabolic pathways and non-ER “response to stress” network, whereas “protein processing in ER” links mostly only to “protein catabolic process”. Genes circled in red indicate known rate-limiting metabolic enzymes.

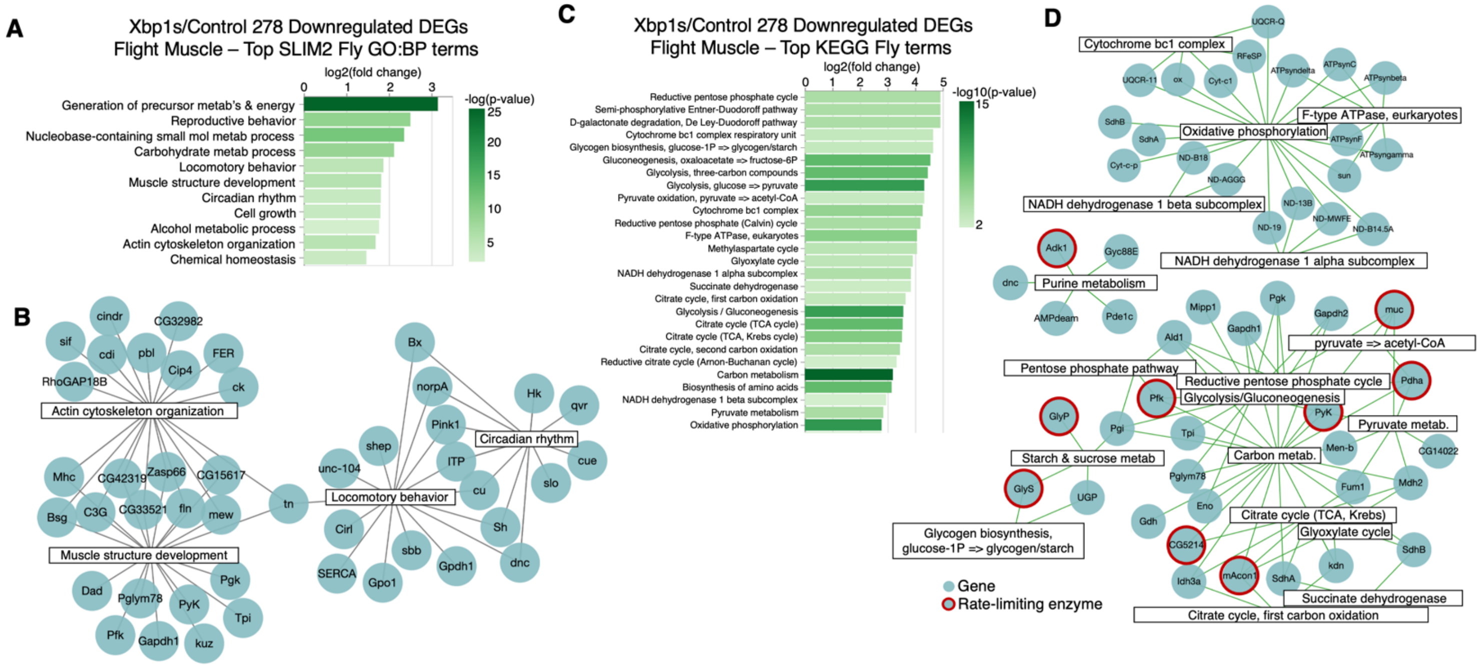


**Supplemental Figure 5. Gene Ontology (GO) and KEGG enrichment of downregulated differentially expressed genes (DEGs) in Xbp1-activated flight muscle. A)** PAthway, Network, and Gene-set Enrichment Analysis (PANGEA) overrepresentation analysis of *Drosophila* SLIM2 GO Biological Process (BP) terms for all downregulated DEGs (278 genes total) in flight muscle. Top GO BP term enrichment is shown. **B)** Gene node plots for select enriched terms related to muscle function in Xbp1s flight muscle downregulated DEGs. Each gene/node represents a significantly downregulated DEG. **C)** Overrepresentation analysis showing most enriched *Drosophila* KEGG terms in downregulated flight muscle DEGs. **D)** Gene node plots for select enriched terms related to oxidative phosphorylation, carbohydrate metabolism, and purine metabolism in Xbp1s flight muscle downregulated DEGs. Genes circled in red indicate known rate-limiting metabolic enzymes.

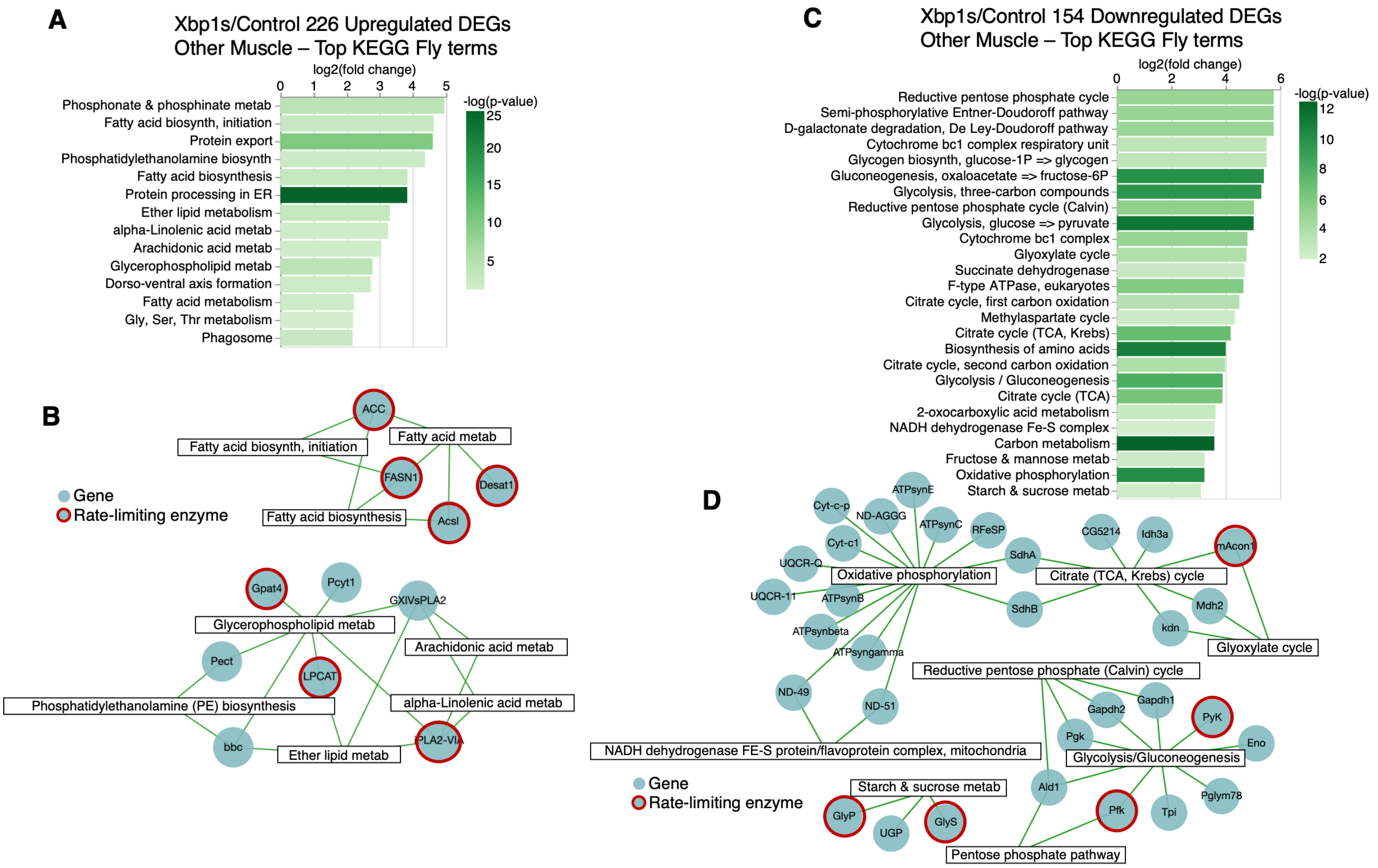


**Supplemental Figure 6. KEGG enrichment of upregulated and downregulated differentially expressed genes (DEGs) in Xbp1-activated "other muscle". A)** PAthway, Network, and Gene-set Enrichment Analysis (PANGEA) overrepresentation analysis of *Drosophila* KEGG terms for all upregulated DEGs (226 genes total) in other muscle. Top KEGG term enrichment is shown. **B)** Gene node plots for select enriched terms in Xbp1s other muscle upregulated DEGs. Each gene/node represents a significantly upregulated DEG. Similar to flight muscle, KEGG metabolic terms related to lipid synthesis were significantly enriched. **C)** Overrepresentation analysis showing most enriched *Drosophila* KEGG terms in downregulated other muscle DEGs (154 genes total). **D)** Gene node plots for select enriched terms in Xbp1s other muscle downregulated DEGs. Similar to flight muscle, oxidative phosphorylation and carbohydrate metabolism terms were significantly enriched. Genes circled in red indicate known rate-limiting metabolic enzymes.

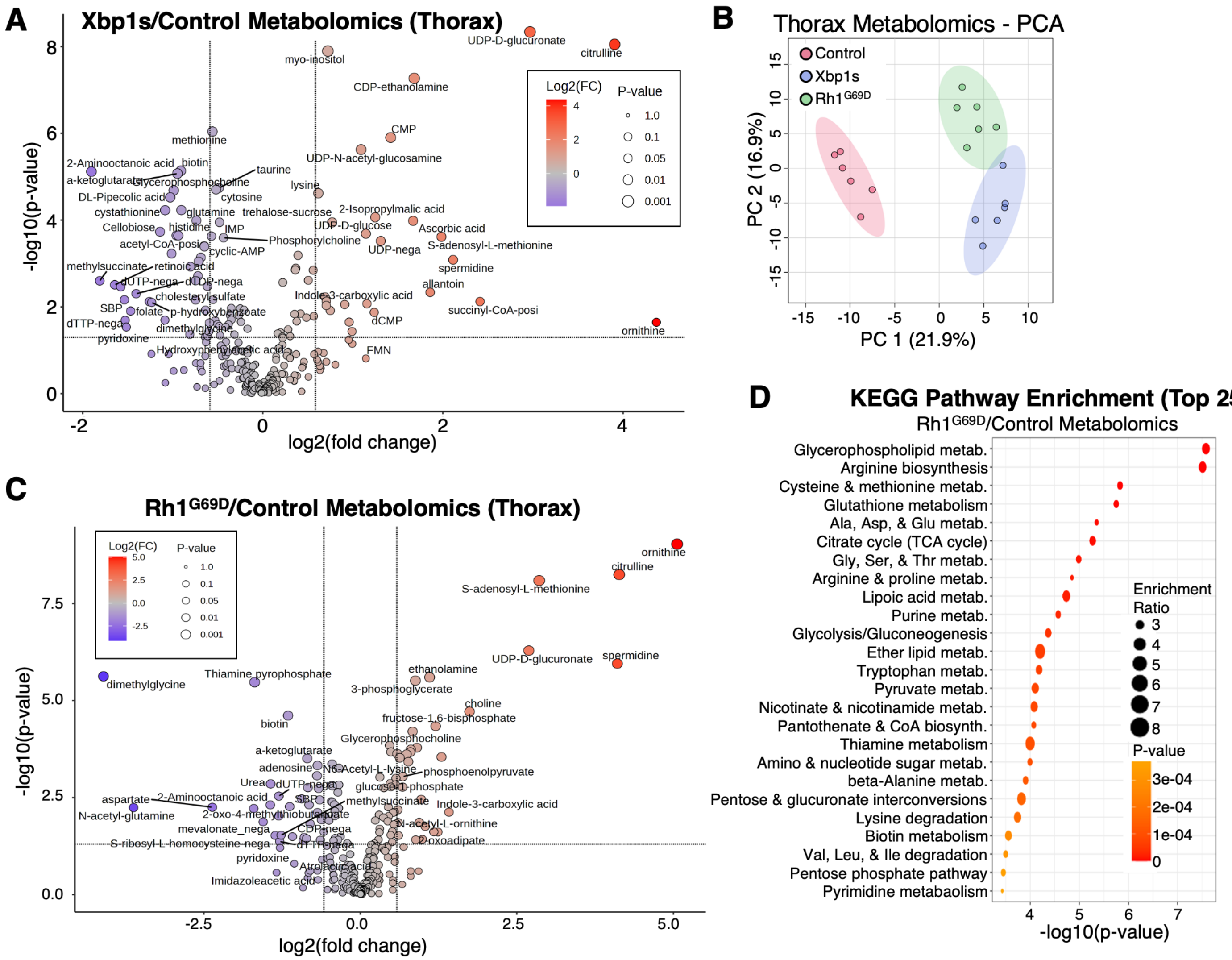


**Supplemental Figure 7. Characterization of targeted polar metabolomics from 6d UPR-activated thoraces. A)** Volcano plot of metabolite abundances between control and Xbp1s-expressing thoraces, averaged across 6 biological reps each condition, 1 rep = 25 thoraces. **B)** Principal component analysis (PCA) of all thorax samples. **C)** Volcano plot of metabolite abundances between control and Rh1$^{G69D}$-expressing thoraces, averaged across 6 biological reps each condition, 1 rep = 25 thoraces. **D)** Most enriched annotated KEGG terms in list of changed polar metabolites from Rh1$^{G69D}$ versus control thoraces. Enrichment indicates pathway disruption compared to control, not necessarily increased or decreased abundance.

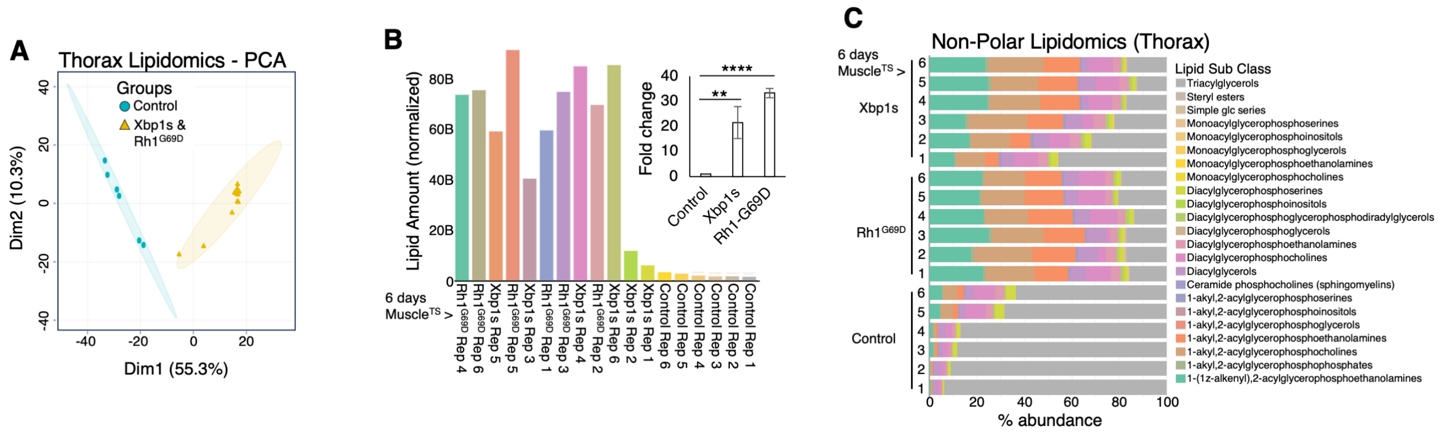


**Supplemental Figure 8. Characterization of untargeted nonpolar lipidomics from 6d UPR-activated thoraces. A)** Principal component analysis (PCA) of all thorax samples. Xbp1s- and Rh1$^{G69D}$-expressing thoraces clustered together, indicating little variation between UPR-activated samples. **B)** Bar graphs of normalized total lipid amount across all replicates (1 rep = 25 thoraces). Fold change compared to control shown. Significance was determined using unpaired T-test. **C)** LipidSig2.0 determined lipid sub class composition for each replicate.

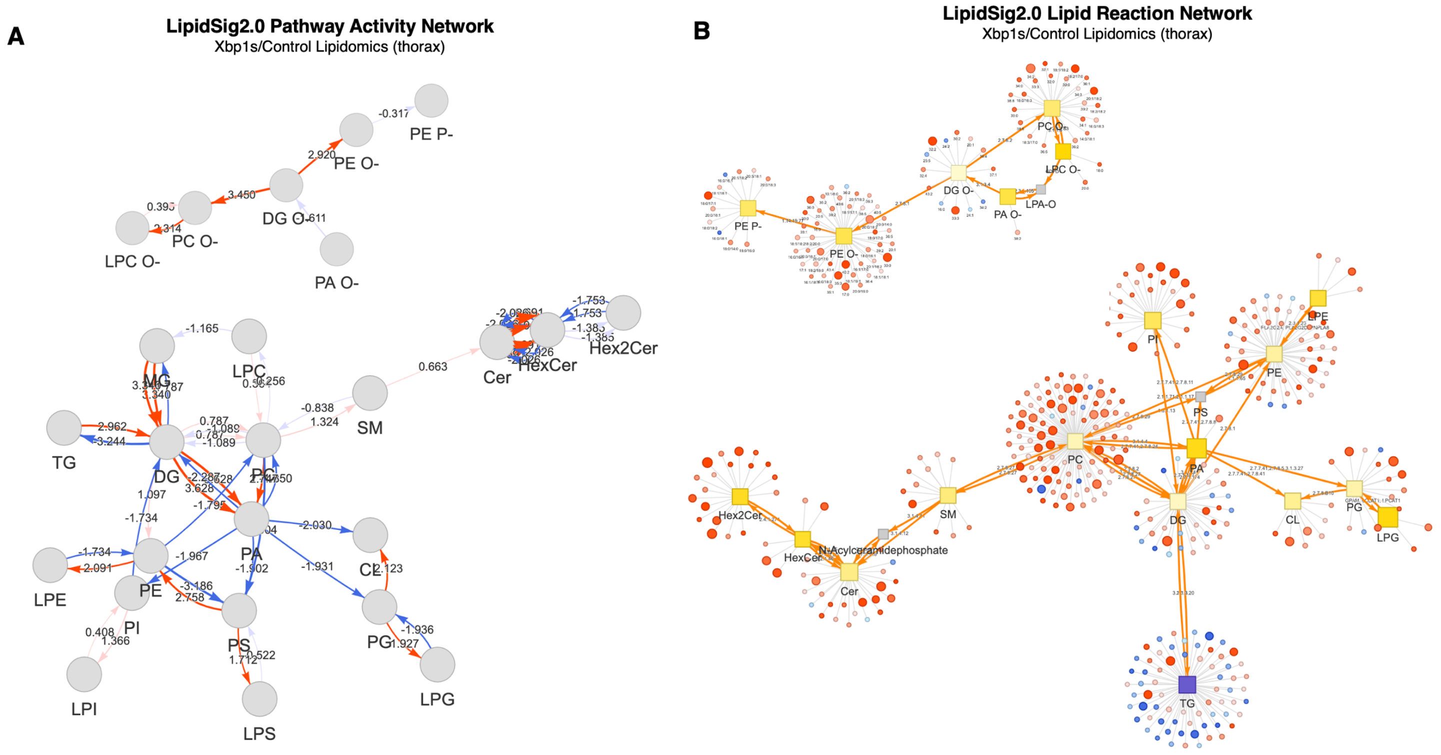


**Supplemental Figure 9. LipidSig2.0 pathway and reaction network analyses for 6d UPR-activated lipidomics. A)** Differentially abundant lipid species were first determined between Xbp1s and control thoraces and then used for pathway activity network analysis. Red arrows indicate predicted upregulated reaction directions, while blue arrows predict downregulated directions. Each node represents a lipid class differentially abundant in Xbp1s thoraces. **B)** Lipid reaction network analysis visualization from differentially abundant lipid species between Xbp1s and control thoraces. Arrows indicate directionality, squares represent lipid classes, and circles surrounding lipid classes represent individual lipid species detected within those classes. Red circles indicate increased abundance of that specific lipid, while blue indicates decreased abundance.

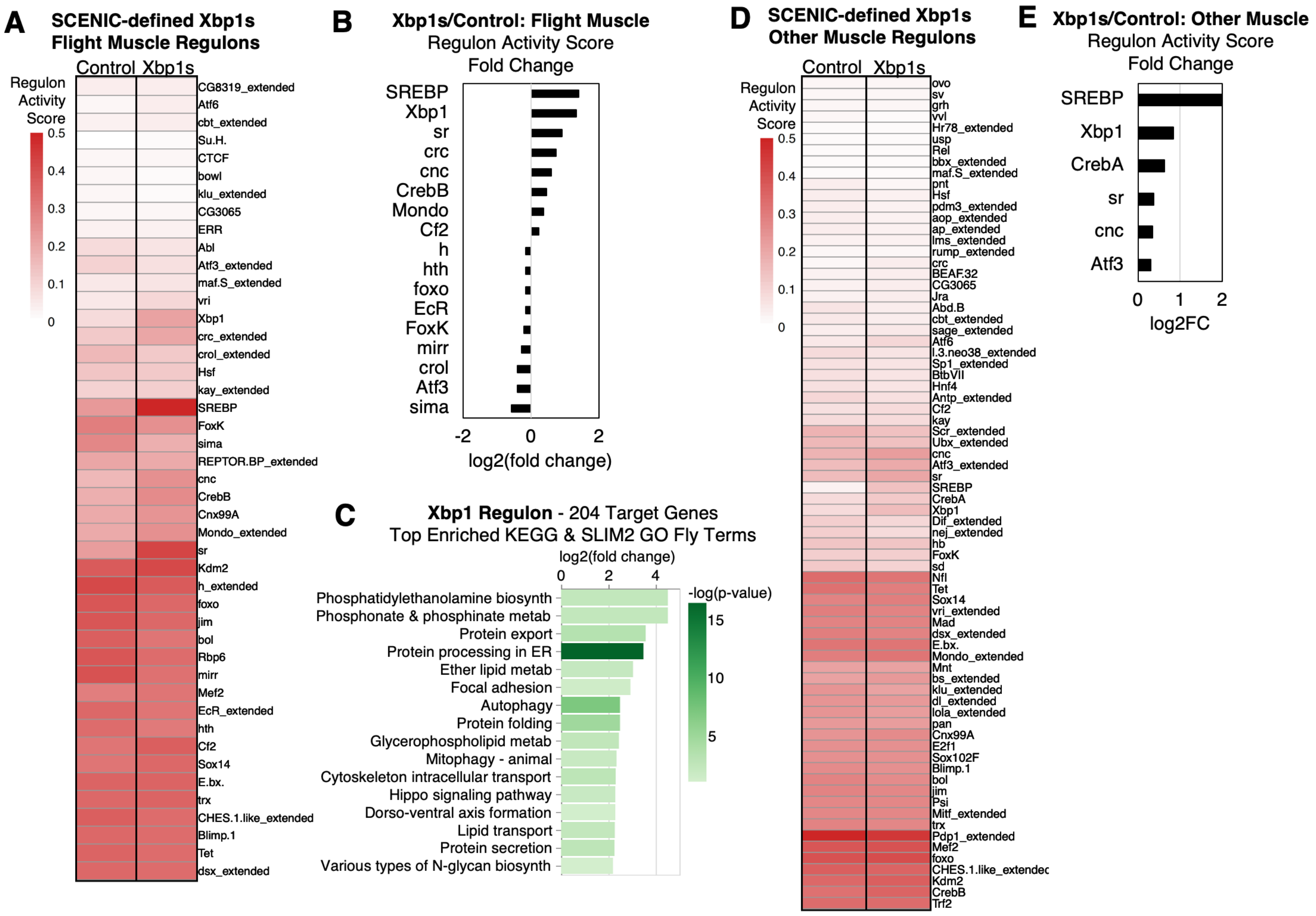


**Supplemental Figure 10. Scoring SCENIC-identified transcription factor (TF) regulons in Xbp1-activated muscle clusters. A)** SCENIC was ran 20 times in Xbp1s flight muscle cells. Regulons and target genes appearing >80% runs were compiled as "consensus" TF regulons and subsequently scored in Xbp1s flight muscle cells using AUCell. To determine changed activity compared to control, these Xbp1s regulons were also scored using AUCell in control flight muscles. Heatmap depicts mean AUCell regulon activity score for control vs Xbp1s flight muscle. **B)** Fold change in Xbp1s/control activity scores were determined from "**A**". Only regulons with activity score >1.0 and fold change >1.2 or <0.8 were kept. **C)** Overrepresentation analysis of enriched KEGG and SLIM2 GO BP fly terms in 204 target genes of Xbp1s-expressing flight muscle Xbp1 regulon. **D-E)** Identical analysis and visualizations as in "**A-B**" but in "other muscle" cells rather than flight muscle. This method did not identify regulons with significantly decreased activity after Xbp1 activation in other muscle cells.

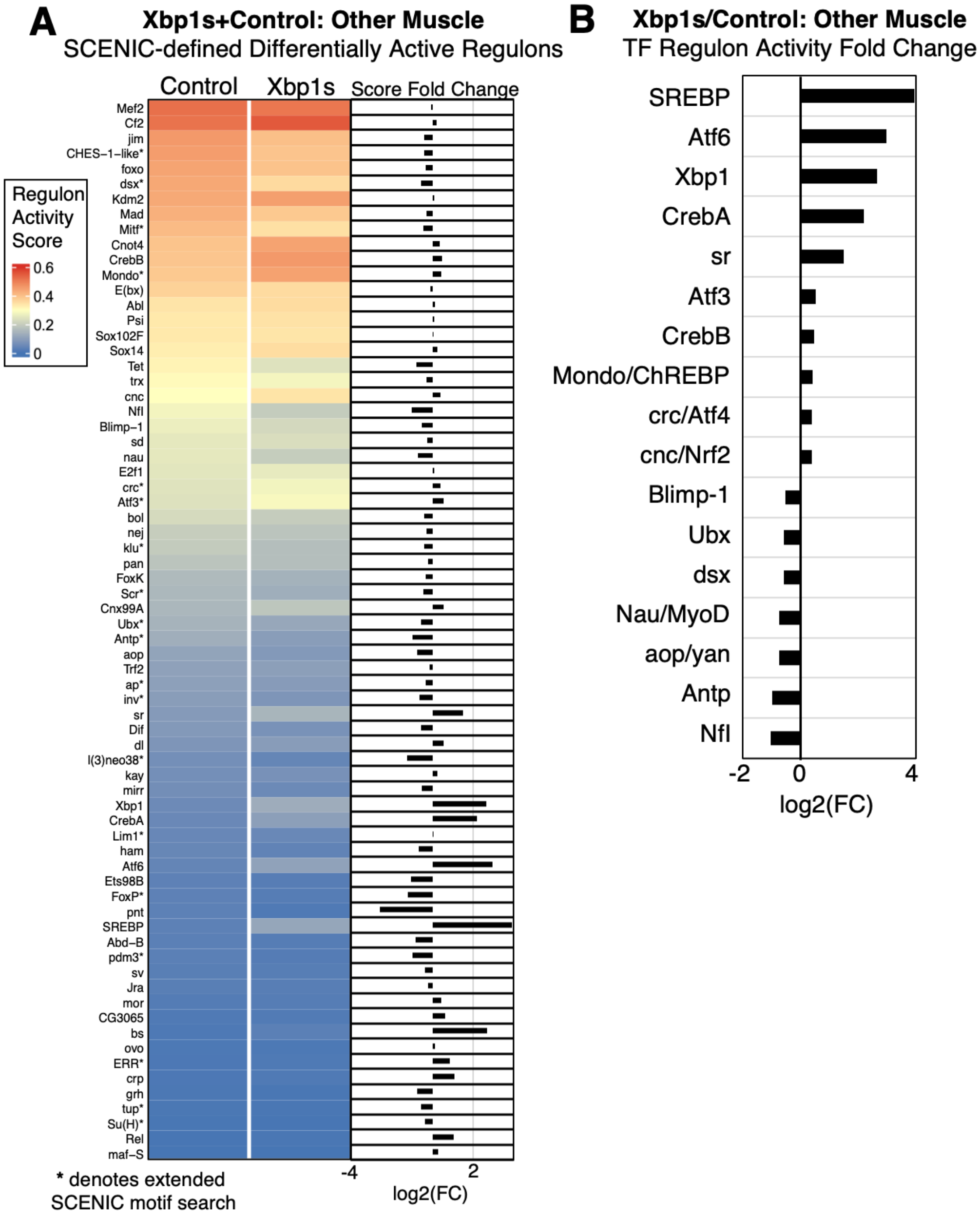


**Supplemental Figure 11. Alternative SCENIC strategy to identify differentially active regulons between control and Xbp1s other muscle cells. A)** Heatmap of AUCell activity scores and fold change between Xbp1s and control other muscle cells of differentially active regulons (i.e., alternative SCENIC strategy using both control and Xbp1s other muscle cells as co-expression matrices input for regulon building). **B)** Only regulons with activity score >1.0 and fold change >1.2 or <0.8 were kept. Compared to using only Xbp1s-generated SCENIC regulons (Supplemental Figure 10), including control other muscle cells increased ability to predict regulons with decreased activity between Xbp1s and control cells.

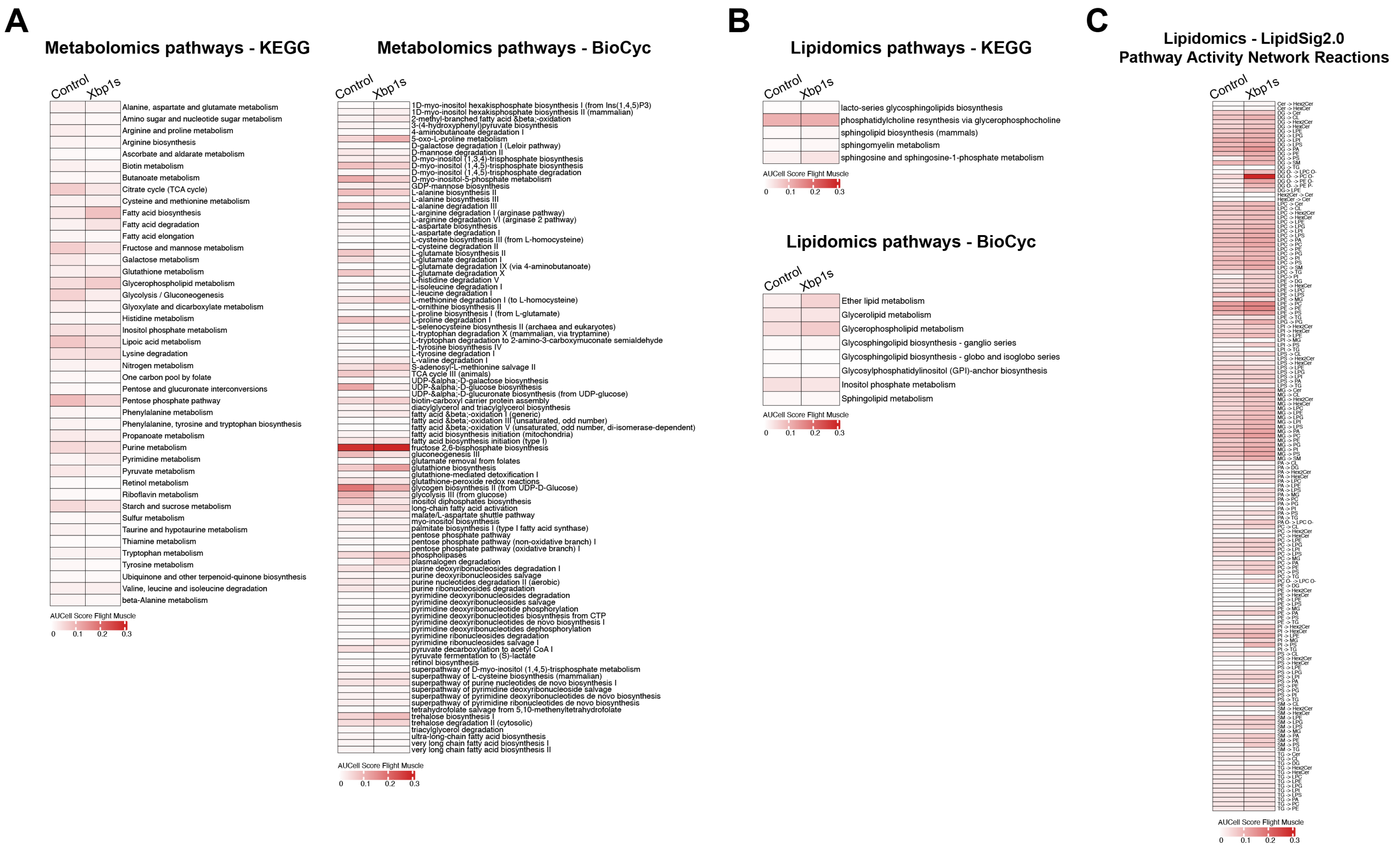

**Supplemental Figure 12. snRNA-seq AUCell scoring of all altered metabolomic and lipidomic pathways in Xbp1-activated flight muscle. A-B)** From significantly altered metabolite (**A**) and lipid (**B**) abundances determined by mass spectrometry metabolomics and lipidomics, we curated all relevant KEGG or BioCyc annotated pathways affected. Using AUCell, expression of annotated genes at the pathway-level were scored in control and Xbp1s snRNA-seq flight muscle to determine network-level changes in transcription. **C)** AUCell scoring in control and Xbp1s flight muscle cells of condensed lipid reactions determined by LipidSig2.0 pathway activity analysis (Supplemental Figure 9A). Fly orthologues were first mapped onto human gene annotations from LipidSig2.0/

**A**

**Upregulated Lipid Metabolism - DEGs**

| Symbol | Gene | Avg_log2FC | p-value |
|---|---|---|---|
| GXIVsPLA2 | FBgn0036545 | 5.361465974 | 2.6089E-206 |
| Spt-I | FBgn0086532 | 4.704367892 | 4.24337E-37 |
| CG7149 | FBgn0031948 | 4.222653347 | 5.92815E-34 |
| iPLA2-VIA | FBgn0036053 | 3.982751791 | 0 |
| Acsl | FBgn0263120 | 3.717672688 | 0 |
| CG9149 | FBgn0035203 | 3.351453075 | 7.39974E-51 |
| CG9706 | FBgn0036662 | 3.090251829 | 1.76849E-35 |
| Pect | FBgn0032482 | 2.746476803 | 1.8655E-187 |
| bbc | FBgn0033844 | 2.660056394 | 3.1659E-277 |
| Pcyt1 | FBgn0041342 | 2.559930515 | 0 |
| PIG-A | FBgn0034270 | 2.351576413 | 4.70292E-05 |
| Sk2 | FBgn0052484 | 2.306546306 | 2.49219E-14 |
| PIG-H | FBgn0037535 | 2.132572167 | 1.26415E-14 |
| PIG-L | FBgn0038763 | 2.122578458 | 3.58563E-14 |
| Dhap-at | FBgn0040212 | 2.072254567 | 1.08372E-15 |
| PIG-N | FBgn0033479 | 2.020089429 | 4.29887E-07 |
| Gpat4 | FBgn0034971 | 1.911725053 | 4.26505E-94 |
| Pld3 | FBgn0032923 | 1.909255522 | 1.95914E-18 |
| GlcT | FBgn0067102 | 1.793258649 | 2.41883E-07 |
| PAPLA1 | FBgn0031990 | 1.688512892 | 3.5447E-138 |
| Pcyt2 | FBgn0035231 | 1.583775623 | 7.94791E-19 |
| CG1941 | FBgn0033214 | 1.578691886 | 1.77921E-05 |
| beta4GalNAcTA | FBgn0027538 | 1.437763796 | 8.10972E-07 |
| Dgkepsilon | FBgn0020930 | 1.428373531 | 5.88576E-11 |
| LPCAT | FBgn0052699 | 1.379990452 | 1.37E-115 |
| Sk1 | FBgn0030300 | 1.285624285 | 0.000923288 |
| PIG-F | FBgn0036893 | 1.275411488 | 1.80854E-12 |
| CG2818 | FBgn0031566 | 1.222834687 | 3.11865E-34 |
| CG12746 | FBgn0037341 | 1.127891957 | 5.80121E-08 |
| CG10425 | FBgn0039304 | 1.011784787 | 4.81409E-06 |
| schlank | FBgn0040918 | 0.979212039 | 2.75014E-13 |
| SMSr | FBgn0052380 | 0.969703116 | 1.98082E-11 |
| CG3961 | FBgn0036821 | 0.923678431 | 2.63619E-41 |
| Pis | FBgn0030670 | 0.922705372 | 6.53214E-18 |
| CG4598 | FBgn0032160 | 0.917741316 | 0.003210233 |
| CG31717 | FBgn0051717 | 0.901656904 | 0.000429875 |
| eas | FBgn0000536 | 0.838985534 | 0.000339806 |
| PGAP5 | FBgn0031997 | 0.826925873 | 0.000730674 |
| ACC | FBgn0033246 | 0.802432349 | 1.1001E-32 |
| ifc | FBgn0001941 | 0.790909252 | 0.007848762 |
| CG18815 | FBgn0042138 | 0.778071602 | 5.78767E-13 |
| FASN1 | FBgn0283427 | 0.60935615 | 2.08542E-14 |
| yip2 | FBgn0040064 | 0.586022054 | 1.65338E-05 |

**B**

**Upregulated Non-Lipid Metabolism - DEGs**

| Symbol | Gene | Avg_log2FC | p-value |
|---|---|---|---|
| CG32196 | FBgn0052196 | 3.3635001 | 2.0593E-19 |
| IP3K2 | FBgn0283680 | 3.03703144 | 2.9283E-10 |
| Inos | FBgn0025885 | 2.73680057 | 5.0844E-91 |
| l(2)k01209 | FBgn0022029 | 2.57234861 | 7.59E-116 |
| isoQC | FBgn0036999 | 2.4310914 | 2.248E-09 |
| NTPase | FBgn0024947 | 2.40793589 | 1.079E-286 |
| RnrS | FBgn0011704 | 2.30890869 | 4.6669E-05 |
| GstE11 | FBgn0034354 | 2.24229743 | 3.9972E-48 |
| Papss | FBgn0020389 | 2.02215 | 5.5029E-11 |
| PHGPx | FBgn0035438 | 1.8272007 | 4.0357E-82 |
| Idh | FBgn0001248 | 1.69801799 | 3.5657E-71 |
| CG1673 | FBgn0030482 | 1.69486103 | 1.2997E-40 |
| SpdS | FBgn0037723 | 1.69211399 | 1.9666E-09 |
| CG10365 | FBgn0039109 | 1.65602088 | 0 |
| Ldh | FBgn0001258 | 1.63556486 | 6.6798E-12 |
| Mccc2 | FBgn0042083 | 1.62209406 | 3.5119E-11 |
| Dak1 | FBgn0028833 | 1.42265358 | 1.269E-15 |
| dnk | FBgn0022338 | 1.36246457 | 0.00014843 |
| GstD9 | FBgn0038020 | 1.30161657 | 1.6492E-15 |
| l(3)72Dp | FBgn0263607 | 1.29855664 | 2.6555E-07 |
| GstD3 | FBgn0010039 | 1.22120053 | 7.5556E-05 |
| Gclc | FBgn0040319 | 1.15917936 | 1.7497E-63 |
| CG5276 | FBgn0037900 | 1.15680992 | 4.7086E-09 |
| Eip55E | FBgn0000566 | 1.05463501 | 3.4817E-05 |
| CG5171 | FBgn0031907 | 1.04135487 | 4.3035E-45 |
| GstD1 | FBgn0001149 | 0.98698161 | 3.767E-89 |
| pyd3 | FBgn0037513 | 0.96106889 | 3.9898E-06 |
| l(3)72Dr | FBgn0263608 | 0.91896546 | 0.00195166 |
| Uck | FBgn0263398 | 0.87856289 | 1.2528E-08 |
| Dhod | FBgn0000447 | 0.84744959 | 0.00031426 |
| ACC | FBgn0033246 | 0.80243235 | 1.1001E-32 |
| r-l | FBgn0003257 | 0.72962161 | 1.357E-09 |
| sprt | FBgn0082585 | 0.66348092 | 1.8305E-09 |
| CG8360 | FBgn0032001 | 0.65460595 | 3.8867E-07 |
| Sam-S | FBgn0005278 | 0.65417131 | 4.6031E-28 |
| Ipk1 | FBgn0050295 | 0.63411372 | 8.0069E-05 |
| yip2 | FBgn0040064 | 0.58602205 | 1.6534E-05 |
| CRMP | FBgn0023023 | 0.57588158 | 0.01169486 |
| Treh | FBgn0003748 | 0.52550866 | 1.4225E-43 |

**C**

**Downregulated Non-Lipid Metabolism - DEGs**

| Symbol | Gene | Avg_log2FC | p-value |
|---|---|---|---|
| CG17224 | FBgn0031489 | -6.14337 | 2.8858E-14 |
| CG11052 | FBgn0040524 | -4.3344217 | 4.385E-27 |
| Argk | FBgn0000116 | -3.6405934 | 0 |
| CG3635 | FBgn0032981 | -3.3752048 | 3.9261E-16 |
| Pde6 | FBgn0038237 | -3.3082585 | 4.8096E-12 |
| Pde1c | FBgn0264815 | -3.1222153 | 0 |
| kdn | FBgn0261955 | -3.1020483 | 9.147E-235 |
| Gyc88E | FBgn0038295 | -3.0832611 | 1.269E-243 |
| CG6330 | FBgn0039464 | -2.9176781 | 9.9361E-06 |
| Ugt36D1 | FBgn0032713 | -2.820371 | 6.3083E-36 |
| alpha-Est2 | FBgn0015570 | -2.5738863 | 1.5429E-06 |
| Pfk | FBgn0003071 | -2.5669552 | 0 |
| CG42637 | FBgn0261360 | -2.5225276 | 8.9679E-07 |
| UGP | FBgn0035978 | -2.4007168 | 1.492E-130 |
| CG11883 | FBgn0033538 | -2.3859217 | 2.6864E-14 |
| CG14022 | FBgn0031700 | -2.3277033 | 1.9607E-60 |
| CG3842 | FBgn0029866 | -2.2606625 | 7.5581E-10 |
| Gyc76C | FBgn0266136 | -2.2510301 | 1.1019E-06 |
| Tpi | FBgn0086355 | -2.2251194 | 6.6755E-73 |
| norpA | FBgn0262738 | -2.1915616 | 1.941E-210 |
| Gdh | FBgn0001098 | -2.1209732 | 1.0569E-27 |
| Pglym78 | FBgn0014869 | -2.0923487 | 7.0636E-48 |
| PyK | FBgn0267385 | -2.0914822 | 5.0509E-63 |
| Pgm1 | FBgn0003076 | -2.0658565 | 2.198E-23 |
| Adh | FBgn0000055 | -2.0366021 | 2.3402E-10 |
| CG18231 | FBgn0036796 | -1.965323 | 3.6325E-05 |
| Fum1 | FBgn0286222 | -1.9481169 | 2.65E-99 |
| alpha-Est7 | FBgn0015575 | -1.9264037 | 0.00013625 |
| Ady43A | FBgn0026602 | -1.9224965 | 2.9657E-09 |
| GlyS | FBgn0266064 | -1.8538154 | 6.605E-110 |
| Odc1 | FBgn0013307 | -1.8439889 | 0.00215526 |
| AGBE | FBgn0053138 | -1.8168021 | 4.25E-10 |
| GlyP | FBgn0004507 | -1.7843675 | 2.7585E-30 |
| CG17097 | FBgn0265264 | -1.770342 | 2.6948E-42 |
| Mipp1 | FBgn0026061 | -1.7611085 | 1.2045E-37 |
| Adk1 | FBgn0022709 | -1.6844684 | 3.8474E-80 |
| Pgk | FBgn0250906 | -1.6710068 | 3.9344E-35 |
| PH4alphaNE2 | FBgn0039783 | -1.6680448 | 0.01027093 |
| Gal | FBgn0001089 | -1.6672609 | 0.01302846 |
| Eno | FBgn0000579 | -1.6503692 | 3.215E-135 |
| CG15093 | FBgn0034390 | -1.6467263 | 0.00641472 |
| PCB | FBgn0027580 | -1.5927642 | 2.378E-11 |
| CG44243 | FBgn0265178 | -1.5880499 | 0.00867392 |
| SdhB | FBgn0014028 | -1.5461913 | 4.7024E-20 |
| mdy | FBgn0004797 | -1.5388787 | 0.00080233 |

| Symbol | Gene | Avg_log2FC | p-value |
|---|---|---|---|
| CG18284 | FBgn0043825 | -1.5222165 | 4.1459E-11 |
| Gapdh1 | FBgn0001091 | -1.4737472 | 1.3478E-80 |
| CG5214 | FBgn0037891 | -1.466541 | 2.8461E-21 |
| Mdh2 | FBgn0262559 | -1.4642392 | 3.7099E-20 |
| alpha-Est4 | FBgn0015572 | -1.4485845 | 0.01464549 |
| Gld | FBgn0001112 | -1.4423874 | 0.00103686 |
| Idh3a | FBgn0027291 | -1.4386524 | 1.5273E-26 |
| CG9485 | FBgn0034618 | -1.4376291 | 1.3443E-10 |
| CG3483 | FBgn0035005 | -1.4364494 | 0.0119157 |
| Pepck2 | FBgn0034356 | -1.4068629 | 3.3511E-14 |
| CG18258 | FBgn0265267 | -1.4042692 | 0.02521596 |
| CG4829 | FBgn0030796 | -1.3965088 | 0.03087837 |
| mag | FBgn0036996 | -1.3882025 | 0.00091481 |
| Men-b | FBgn0029155 | -1.3875169 | 4.2799E-23 |
| Ald1 | FBgn0000064 | -1.3723312 | 0 |
| CG15547 | FBgn0039809 | -1.361674 | 6.423E-05 |
| SdhA | FBgn0261439 | -1.3340742 | 1.912E-23 |
| CG4546 | FBgn0038373 | -1.3251769 | 0.00024264 |
| CG11598 | FBgn0038067 | -1.3226126 | 0.00717771 |
| dnc | FBgn0000479 | -1.3139099 | 2.378E-110 |
| Got2 | FBgn0001125 | -1.3110942 | 6.6579E-05 |
| CG3809 | FBgn0037995 | -1.3108607 | 0.00847381 |
| CG32549 | FBgn0052549 | -1.3081093 | 4.874E-11 |
| Gapdh2 | FBgn0001092 | -1.2936749 | 8.643E-17 |
| CG30016 | FBgn0050016 | -1.2921719 | 0.00835296 |
| CG17010 | FBgn0032424 | -1.2825861 | 0.01131715 |
| Pdha | FBgn0028325 | -1.2384153 | 9.7633E-17 |
| CG5599 | FBgn0030612 | -1.2268831 | 0.00019028 |
| muc | FBgn0283658 | -1.215143 | 4.4404E-13 |
| CG7069 | FBgn0038952 | -1.2088084 | 0.00931827 |
| CG31872 | FBgn0051872 | -1.1855274 | 5.7869E-11 |
| CG32026 | FBgn0052026 | -1.1757361 | 0.00118838 |
| CG32445 | FBgn0052445 | -1.1626376 | 0.00069102 |
| l(1)G0196 | FBgn0027279 | -1.1314494 | 4.2449E-26 |
| Adk2 | FBgn0283494 | -1.1277729 | 0.04390611 |
| Est-6 | FBgn0000592 | -1.1100617 | 9.7859E-08 |
| Nc73EF | FBgn0010352 | -1.0917749 | 9.079E-230 |
| Pgi | FBgn0003074 | -1.0914228 | 2.4765E-15 |
| Gfat1 | FBgn0287209 | -1.0862503 | 0.02679255 |
| ras | FBgn0003204 | -1.0854568 | 1.1633E-08 |
| Ac13E | FBgn0022710 | -1.0691957 | 0.01158622 |
| b | FBgn0000153 | -1.0626504 | 2.1955E-13 |
| AMPdeam | FBgn0052626 | -1.0581266 | 1.416E-13 |
| mAcon1 | FBgn0010100 | -1.0462477 | 1.9294E-18 |
| Echs1 | FBgn0033879 | -1.030092 | 8.7513E-06 |

**Supplemental Figure 13. All differentially expressed genes (DEGs) between Xbp1s and control flight muscle within altered broad metabolic pathway categories. A-C)** List of all DEGs, sorted by average log2FoldChange in Xbp1s vs control flight muscle cells, identified within broad metabolic categories that were transcriptionally altered at the pathway-level. DEGs listed for upregulated lipid metabolic pathways (**A**), upregulated non-lipid metabolic pathways (**B**), and downregulated non-lipid metabolic pathways (**C**). These DEGs were then intersected with SCENIC- and hdWGCNA-defined regulon target genes (Figure 5) to predict transcription factors that may control these metabolic networks.

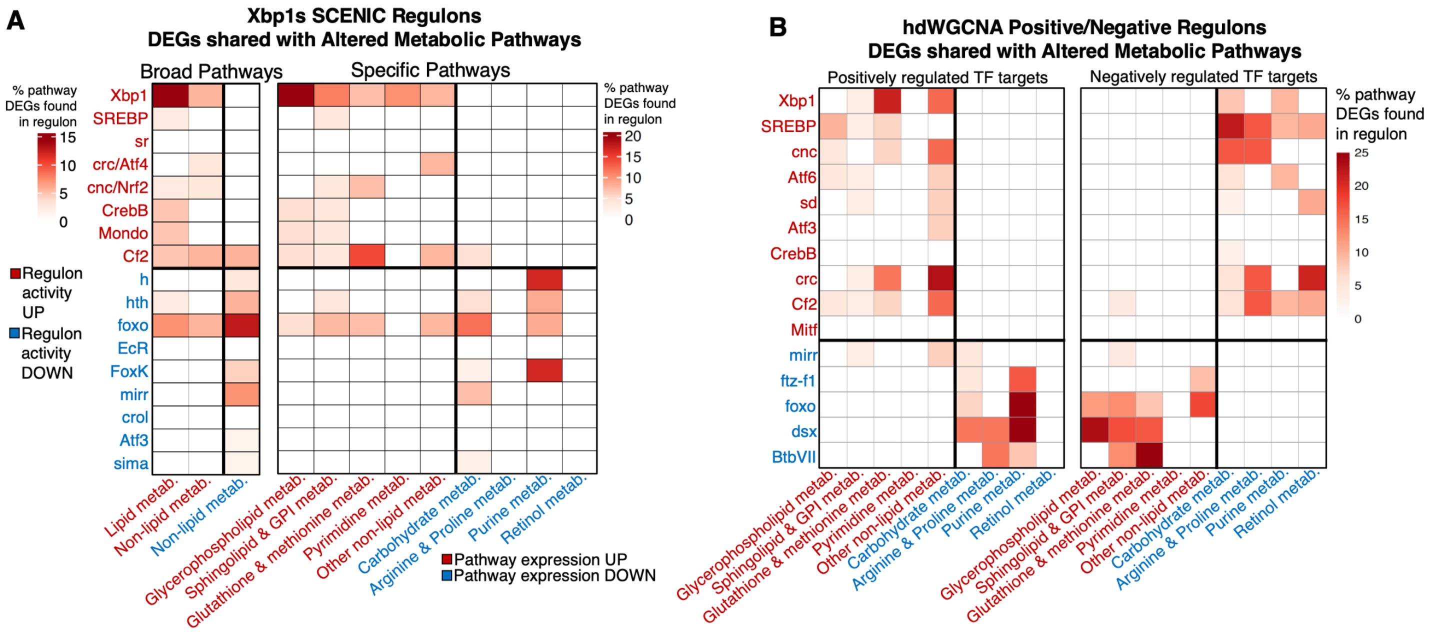


**Supplemental Figure 14. Identifying metabolomic and lipidomic differentially expressed genes (DEGs) within SCENIC and hdWGCNA transcription factor (TF) regulons. A)** Heatmap depicting the percentage of DEGs in a given pathway (categorized into either broad pathways or specific pathways) that were also found in a given SCENIC regulon. While **Figure 5** shows overlap with differentially active regulons (i.e., regulons constructed with control and Xbp1s cells together as input), this heatmap depicts overlap with SCENIC regulons constructed using only Xbp1s-expressing flight muscle cells. Regulons with increased activity (in red) were enriched for DEGs found in upregulated lipid and non-lipid metabolic pathways (in red). Similarly, regulons with decreased activity (in blue) were enriched for DEGs found in downregulated non-lipid metabolic pathways (in blue), with the exception of FoxO, which was predicted to additionally control upregulated metabolic pathways as well. **B)** Same percent DEG analysis of specific metabolic pathways as in “**A**” but with hdWGCNA regulons divided into predicted positively regulated or negatively regulated target genes per TF. Only regulons shared between SCENIC and hdWGCNA are shown.

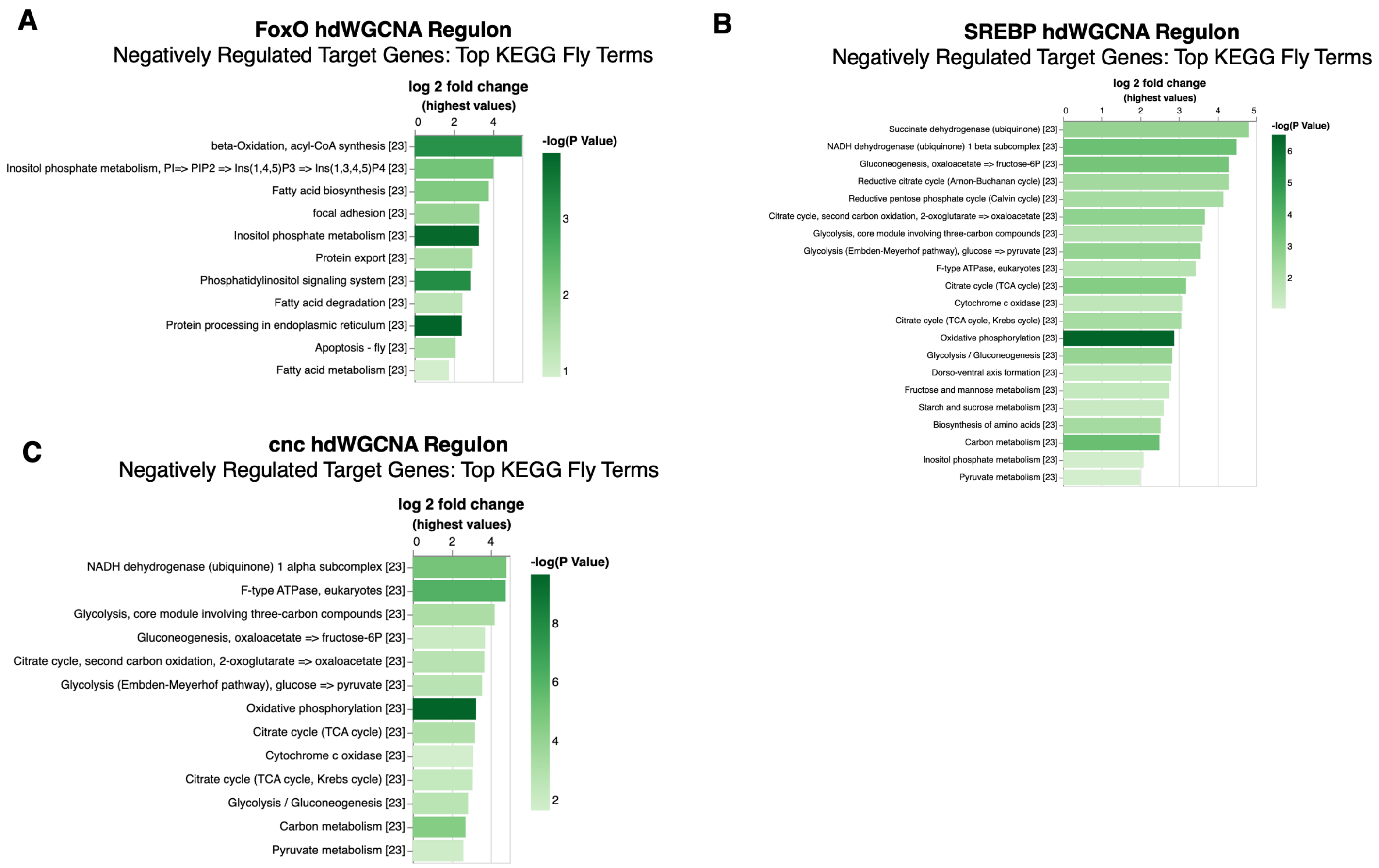


**Supplemental Figure 15. KEGG pathway enrichment of target genes predicted to be negatively regulated by FoxO, SREBP, or cnc. A-C)** Overrepresentation analysis of top enriched KEGG fly terms for all hdWGCNA-predicted genes negatively regulated by FoxO (**A**), negatively regulated by SREBP (**B**), or negatively regulated by cnc/Nrf2 (**C**) in Xbp1-activated flight muscle. FoxO negative targets are enriched for fatty acid metabolism and beta-oxidation. SREBP and cnc/Nrf2 negative targets are enriched for carbohydrate metabolism and oxidative phosphorylation gene sets.
